# A mathematical model for bleb expansion clarifies a role for TalA in regulating blebbing

**DOI:** 10.1101/2025.02.28.640705

**Authors:** Sobana Handi Dinuka Sewwandi de Silva, Emily Movsumova, Jessica Reznik, Zully Santiago, Emmanuel Asante-Asamani, Derrick Brazill

**Affiliations:** Department of Mathematics, Clarkson University, Potsdam, NY 13699, USA; Department of Biological Sciences, Hunter College, New York, NY 10065, USA; Department of Natural Sciences, Baruch College, New York, NY 10010, USA; Academic Affairs, York College, Jamaica, NY 11451, USA; Department of Mathematics, University of Kelaniya, Kelaniya, Sri Lanka

## Abstract

Eukaryotic cells, such as cancer and immune cells, migrate using either pressure-driven blebs or actin polymerization driven pseudopods, with cells preferring to bleb in confined environments where high protrusion forces are required for movement. Blebbing involves a separation of the cell membrane from the cortex, via the detachment of membrane-to-cortex linker proteins. The detached membrane then expands and stabilizes into a spherical cap as a new cortex is formed beneath the protruded membrane while the old one is completely degraded. The role of linker proteins in blebbing has mostly been associated with directing blebs to the leading edge of the cell, where linker enrichment is low, suggesting that cells devoid of linker proteins will bleb profusely. However, experimental work in this study involving *talA* null chemotaxing *Dictyostelium discoideum* cells shows the opposite effect, pointing to an alternative role for TalA. Our quantitative analysis of bleb size and frequency reveals that *talA* null cells produce fewer and smaller blebs in confined environments, pointing to a reduction in their intracellular pressure. A mathematical model of bleb expansion developed and validated with our experimental data supports the hypothesis that linker proteins help the cell maintain intracellular pressure during blebbing by limiting changes to its surface area when pressurized. Our model also identifies elastic and viscous properties of the cell, the assembly rate of the new cortex and disassembly rate of the old cortex as key modulators of change in bleb size induced by weakening the strength of membrane to cortex attachment.

**SIGNIFICANCE:** This work analyzes the role of the membrane-to-cortex linker protein talA in regulating bleb size and frequency during bleb-based chemotaxis. We identified that this protein helps to regulate intracellular pressure by preventing pressure loss due to uniform membrane expansion around the cell. In particular, cells form smaller and less frequent blebs without TalA. Analysis of a mathematical model for bleb expansion demonstrates that weakening the strength of linker proteins is sufficient to reduce the size of blebs. Our model also identifies an important role for actin dynamics and the viscoelastic properties of the cell in regulating the percentage change in blebs due to weakening membrane to cortex attachment. Additionally, we find that cells can partially retract blebs without myosin II by regulating the ratio of polymerization and depolymerization of actin in the reforming cortex.

## INTRODUCTION

Blebs are spherical protrusions that are produced by cells as a mechanism for movement (1–4), particularly as the confinement in the environment increases (5). Bleb-based motility has been observed in many cell types including *Dictyostelium discoideum*, Zebrafish, and some cancer cells. *Dictyostelium* cells constrained under a block of agarose have been observed to move using blebbing (6–8). Blebbing has also been implicated in regulating the migration of *Dictyostelium* cells within a multicellular mound (9). Zebrafish premordial germ cells are able to migrate to the right location in the embryo using blebs (2, 10). Additionally, metastasizing cancer cells undergoing single cell motility have been observed to move with blebs (11–14), especially when drugs reduce the functionality of their actin polymerization machinery (15).

A hallmark of blebbing is the separation of the cell membrane from a local region of the cortex and its subsequent expansion into a spherical cap, powered by prevailing intracellular pressure (4, 16, 17). This pressure is thought to be regulated by cortical tension, generated by myosin II-driven contraction of the acto-myosin cortex (16) or confinement forces within the cellular environment (5). Indeed, myosin deletion cells fail to bleb whereas cells with reduced myosin contractility or confinement produce smaller blebs (5, 18). As the membrane expands, a new cortex reforms beneath the protruded membrane and the old cortex (actin scar), at the original location of the detached membrane, completely degrades to stabilize the bleb (7, 19).

Prior to bleb expansion, the membrane is held to the cortex through membrane-cortex binding proteins (linker proteins). One such protein is talin whose orthologues in *Dictyostelium* are TalA and TalB (20–22). TalA has been observed to cluster at the rear of chemotaxing *Dictyostelium Discoideum* cells (8, 20), where it prefers to bind to stretched actin filaments (23). This rear clustering of linker proteins has been shown to direct blebs to the front of the cell, where very few TalA proteins accumulate. Quantification of experimental observations revealed that the area of the cell with the highest blebbing activity coincides with the area with lowest TalA enrichment (8). In an under-agarose chemotaxis assay, *Dictyostelium* cells devoid of both TalA and TalB were observed to nucleate blebs more frequently (compared to wild type) at random locations around the cell boundary (8). Taken together, these observations led to the conclusion that TalA enrichment is inversely proportional to blebbing and that cells without TalA will bleb profusely.

In a recent study, TalB proteins were found to localize to the leading edge of chemotaxing *Dictyostelium* cells, where actin filaments are not as stretched as those found in the posterior of the cell (23). Phenotypic differences have been observed in single deletions of *talA* and *talB*. For example, *talA* null cells are able to develop normally, however *talB* nulls arrest at the mound stage and cannot form tips (24). On the other hand, *talA* null cells are defective in cytokinesis and forming adhesions with the substrate (21). The distinct localization of these proteins coupled with their phenotypic differences suggests that the two proteins could have distinct roles in regulating blebbing. Additionally, the evidence provided to support the conclusion from (8) that *talA*/*talB* null cells bleb more profusely was more qualitative than quantitative. Taken together, it is unclear whether the observation of more frequent blebbing in the double talin mutant in (8) will be observed in a single deletion of TalA. Additionally, whereas blebbing is well known to involve a reformation of a cortex beneath the protruded membrane, and a degradation of the old cortex, very little work has been done to investigate the role of actin dynamics in the regulation of bleb size.

Recently, mathematical models have help to provided useful insight into the mechanics of cell migration, allowing for the interrogation of questions that may be beyond the current scope of experimentation (25). For instance, boundary energy-based models have helped to clarify how bleb nucleation sites are determined (8, 19), geometric models have helped to characterize bleb shape (26) while fluid mechanics based models have shed light on the dynamics of intracellular pressure gradients during multiple blebbing events (27–29). Our work examines the role of TalA in regulating blebbing during confined chemotaxis. Contrary to the existing school of thought, we find that *talA* null *Dictyostelium* cells, chemotaxing towards cyclic AMP under agarose, generate significantly fewer blebs than wild type cells. Using a combination of mathematical modeling and quantification of bleb size and cell area from experiments, we conclude that TalA helps to maintain the intracellular pressure necessary for bleb formation, so that in its absence the driving pressure to create blebs is reduced. By simulating our mathematical model under various parameters we are able to explore how blebbing cells with different viscoelastic properties as well as actin dynamics may respond to weakening the strength of membrane to cortex attachment.

## MATERIALS AND METHODS

### Strain and culture conditions

The *Dictyostelium discoideum* cells were grown in shaking culture in HL5 medium with glucose (ForMedium), 100 IU/mL penicillin, and 100*μg*/*mL* streptomycin (Amresco) at 150 rpm at 22^0^𝐶 or on 100mm culture plates. *talA* null (DBS0236180) cells were ordered from Dictybase.org and grown on culture dishes. To generate the GFP expressing lines, both Ax2 (wildtype) and *talA* null lines were electroporated with the LifeAct-GFP plasmid, a generous gift from Dr. Chang-Hoon Choi (30), and selected with G418 (4-20 μ𝑔/*mL*). For details on transformation protocal, clone selection and confirmation of cell lines see (31). The cell lines were sub-confluent in log phase (1 × 10^6^*cells*/*mL* to 4 × 10^6^*cells*/*mL*) from shaking prior to use for experiments (32).

### Cyclic-AMP (cAMP) under agarose assay

A preheated (90^0^𝐶, 1 *min*) number 1 German borosilicate sterile 2 well chambered coverglass slide (ThermoFisher) was loaded with 750μ𝐿 of melted 0.4% or 0.7% Omnipur agarose (EMD Millipore) laced with 1mg/mL of 70,000 MW Rhodamine B isothiocyanate-dextran (Sigma-Aldrich). Once solidified, half of the gel was removed from the well and the remaining portion was slid across to the middle of the chamber, creating two wells with one on each side of the gel. To create the cAMP gradient, 4μ𝑀 cAMP was loaded into one well and incubated for 40 minutes. 1 × 10^5^ to 2 × 10^5^ cells competent for cAMP chemotaxis were loaded into the other well and allowed to settle and crawl under the agarose for 30 minutes prior to imaging. See (32) for additional details.

### Live imaging and microscopy

All imaging data was acquired using a Leica DMI-4000B inverted microscope (Leica Microsystems Inc.) mounted on a TMC isolation platform (Technical Manufacturing Corporation) with a Yokogawa CSU 10 spinning disc head and Hamamatsu C9100-13 ImagEM EMCCD camera (Perkin Elmer) with diode lasers of 491 nm, 561nm, and 638 nm (Spectra Services Inc.) [104]. LifeAct-GFP and RITC-dextran were excited using the 491nm and 561nm lasers, respectively. Images were taken over the course of 30 seconds a 100x/1.44 oil immersion objective at maximum camera speed with exposure times of 0.800 seconds for GFP and 0.122 seconds for RITC channels, resulting in intervals of 1.66 seconds. Images were acquired using Volocity 5.3.3 (Perkin-Elmer) and processed in ImageJ. Cell motility structures with respect to their membrane positions were captured in real-time.

### Image processing

As cells crawled under a RITC-dextran laced agarose gel, actin structures were visualized by LifeAct-GFP while membrane position was visualized as a negative against the RITC background. The open access software ImageJ (https://imagej.nih. gov/ij/) was used. The raw data for each series, consisting of the membrane and cortex channels, were split and separated from each other and adjusted for optimal brightness and contrast using ImageJ. Once complete, the two channels (membrane and cortex) were merged together with the cortex in green and the membrane in red. The merged images were subsequently saved as an .avi file without any compression to be used for detection of blebs and calculation of bleb area. Note that, due to the confinement imposed by the agarose gel, cells flatten and adopt a ’pancake’ shape with a near uniform cross sectional area. Thus our area measurements can be translated into cell/bleb volume by multiplying by the height of the gap between the agarose gel and the slide.

### Determining bleb frequency, bleb area and cell area

We analyzed the first 20 frames to determine the bleb frequency and area, since there were noticeable qualitative effects of the lasers on the cell behavior afterwards. To detect blebs, we looked for a clear separation of the membrane (from RITC-dextran channel) and a reformation of the cortex (from LifeAct-GFP channel). The number of blebs generated by a cell within the first 20 frames of observation was recorded as the frequency. The bleb area was calculated from the stabilized bleb, with a fully reformed bleb cortex and degraded actin scar (old cortex). The bleb area was calculated using our in-house Bleb Detection Algorithm (19). This significantly cut down on the time required to process all cells. The algorithm takes as input a sequence of cell boundary coordinates (extracted from the cortex images) and returns the number and area of blebs detected within the time duration as well as the cell area. The number of blebs detected was compared with our manual counts. For any missed blebs, usually due to small size, the area was calculated manually using ImageJ. The polygonal tool in ImageJ was used to create an outline of the bleb. The resulting boundary was smoothed using the spline fit tool. Area measurements were obtained from the Analyze menu in ImageJ. A similar manual process was used to calculate the cell area.

### Statistical analysis

Bleb and cell area as well as bleb frequency measurements were obtained from two independent experiments for each confinement condition. Box plots were created to visualize the distribution of the bleb frequency and area. We pooled together the data from all trials under the same pressure condition, since they had similar distribution. The Mann Whitney U test was used to check for significant differences between groups. Statistical analysis was done in R-studio.

### Theoretical modeling

Bleb-based migration occurs in three-dimensional space with cells often modeled as spheres and blebs as hemispherical protrusions. In our experiments, cells crawl on the surface of a two-dimensional slide under the weight of an agarose gel, which creates a pseudo three-dimensional environment. Under such confinement, cells adopt a ‘pancake’ shape with an almost uniform vertical cross section. In this context, we simplify the cell geometry to two-dimensions by considering change in the cell shape during blebbing on a fixed horizontal plane.

We model bleb expansion in two stages. First, a simple one dimension model (1D) is developed to track the boundary displacement along a radial line drawn from the center of the cell through a single point on the boundary, as illustrated in Fig. 1A bottom. This is intended to mimic the separation distance between the bleb boundary and the cell boundary at any time. Next we extend this model to two-dimensions by tracking displacement along a continuous distribution of radial lines through the cell boundary, as illustrated in Fig. 1B. This permits a more complete description of the dynamics of the cell boundary and cytoplasm as well as forces during blebbing, with the 1D model as a fundamental building block. We present a description of both models here and defer detailed derivations to the supplemental methods.

**Figure 1:**
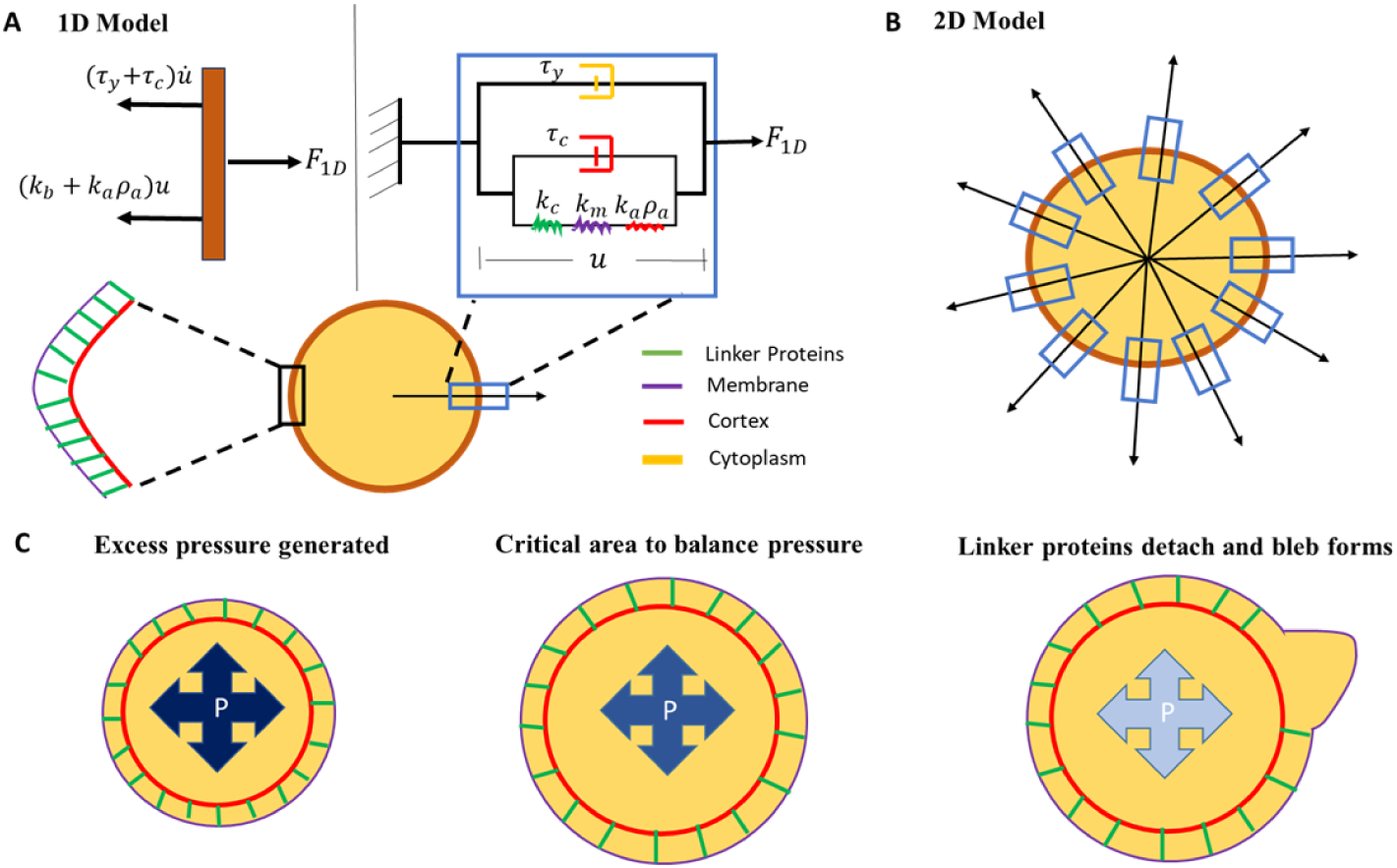
Mechanical assumptions for mathematical model. A) 1D mechanical element of the cell. B) 2D organization of 1D mechanical elements. C) Illustration of bleb initiation process. Lighter color denotes lower pressure, 𝑃.

### 1D dynamic model for bleb expansion

During blebbing, the cell boundary extends due to motion of cytoplasmic fluid. Our primary assumption about the cytoplasm is that it is a viscous fluid with viscosity coefficient 𝜏_*y*_ (*nNs*/*μm*^3^). This assumption has been used in other bleb expansion models and found to yield biologically relevant bleb dynamics (28). We consider the cell boundary as a composite body comprising of an outer membrane which is supported by an inner cortex with linker proteins connecting the two structures. The membrane is a phospholipid bilayer, modeled as an elastic material with elastic stiffness *k_m_* (*nN*/μ*m*^3^). The dense network of crosslinking and motor proteins holding branched actin filaments together in the cortex give it elastic and viscous properties, with the viscous behavior stemming from the dynamic renewal of actin filaments through polymerization and depolymerization (33, 34). Thus, the cortex can be considered as a Kelvin-Voigt viscoelastic solid, with viscosity coefficient 𝜏_𝑐_ (*nNs*/μ*m*^3^) and elastic stiffness 𝑘_𝑐_ (*nN*/μ*m*^3^). Proteins linking the membrane to the cortex undergo continuous binding and unbinding, with a constant binding rate *k_on_* (*S*^−1^) and a force dependent unbinding rate *k_off_* (*S*^−1^). At any point along the boundary, the local density of bound linkers, 𝜌_𝑎_ (μ*m*^−2^) is assumed to induce an elastic resistance of 𝑘_𝑎_ 𝜌_𝑎_ to membrane separation with stiffness 𝑘_𝑎_ (*nN*/μ*m*) per linker protein.

Since the motion of the cell boundary is driven by the flow of the cytoplasm, we assume that these two components (cytoplasm and boundary) experience the same velocity 𝑢· (μ*m*/*S*) with displacement 𝑢(μ*m*). Note that the displacement 𝑢 is a scalar quantity in 1D since it is measured along a radial line through the cell boundary, as shown in Fig. 1A bottom. The assumption of a common velocity for cytoplasm and boundary represents a no-slip condition which has been used in recent models of bleb formation (28). We represent the common displacement of the cytoplasm and boundary by arranging their mechanical elements in parallel, as shown in Fig. 1A top right.

Prior to bleb initiation, the linker proteins inhibit separation of the membrane from the cortex. The separation of the membrane from the cortex creates a purely elastic boundary which expands under the force of fluid pressure. Shortly after membrane detachment, a new cortex begins to form under the protruded membrane giving the bleb boundary viscoelastic properties. These observations motivate our treatment of the boundary as a single viscoelastic structure with time-dependent mechanical properties. Its elastic component has stiffness contributions from the membrane (*k*_*m*_), cortex (*k*_*c*_) and linker proteins (*k*_𝑎_ 𝜌_𝑎_), arranged in series as shown in Fig. 1A top right. The viscous component, purely due to the cortex viscosity 𝜏_*c*_, is arranged in parallel with the elastic element. As a new cortex is formed beneath the protruded membrane, the old cortex is degraded. The degradation alters the porosity of the cortex and viscosity of the cytoplasm within the bleb, potentially regulating the entry of large proteins, organelles and macromolecules into the bleb. Thus, mechanically the cytoplasm within the bleb (bleb cytoplasm) acts as a damper with time dependent damping coefficient 𝜏_*y*_ (*t*).

### Submodel for boundary velocity and adhesion density

Let 𝐹_1𝐷_ (*nN*/μ*m*^2^) denote the force density from intracellular fluid pressure that drives the expansion of blebs. Then, by our assumption of a parallel arrangement of the cytoplasm and boundary, the force driving bleb expansion is distributed across the cytoplasm 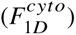 and cell boundary (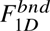) and satisfies the force balance equation

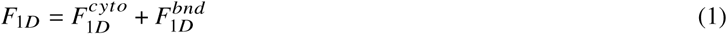

at any point on the cell boundary. The force contribution from the elastic component of the boundary is (*k*_*b*_ (*t*) + *k*_𝑎_ 𝜌_𝑎_)𝑢, where *k*_*b*_ (*t*) is the time-dependent contribution to boundary stiffness from the cortex and membrane (see illustration in Fig. 1A top left). The force from the viscous component of the boundary is proportional to its deformation rate and can be expressed as 𝜏_*c*_ (*t*)𝑢·. Thus, we can write the force across the cell boundary as 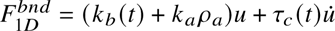. The resistance of the bleb cytoplasm to deformation is proportional to the deformation rate 𝑢· with the force across it modeled as 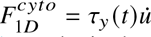 (see illustration in Fig. 1A top left). Combining our forces across the cytoplasm and boundary as shown in Eq. 1 we obtain the force balance

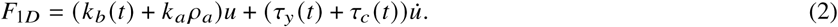

which can be solved for the boundary velocity as

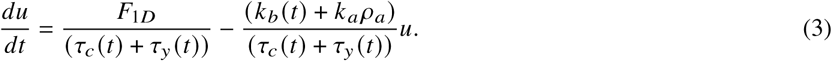

The dynamics of the density of linker proteins, 𝜌_𝑎_ (*t*) is modeled using the classic adhesion model

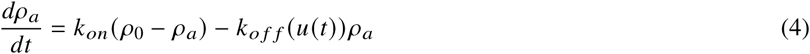

used in Alert’s work in (35) to model membrane peeling during blebbing. Here linker proteins are assumed to bind to the cortex at a rate proportional to the density of unbound linkers 𝜌_0_ − 𝜌_𝑎_ with kinetic constant *k*_𝑜𝑛_ and unbind from the cortex at a rate dependent on the boundary displacement 𝑢, i.e. 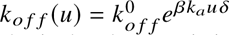. Here, 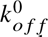 is the rate at which linker proteins detach at zero displacement, 𝛽 is the thermal energy and 𝛿 is the characteristic bond length.

The applied force during bleb expansion 𝐹_1𝐷_ is a pressure gradient force, determined by the difference in fluid pressure between the cytoplasm (high pressure) and the small region of detached membrane at nucleation (low pressure) (36). As the bleb expands, the fluid pressure in the blebbing region increases. The bleb continues to expand until the net pressure between the two regions is zero. In 1D, we we assume this force is initially constant and only varies in time as the boundary expands. We model it as an exponentially decreasing force 𝐹_1𝐷_ (𝑢) = 𝐹_0_𝑒^−*m*𝑢^ with decay rate *m* and initial value 𝐹_0_. This assumption of a constant resting pressure has been used in a recent cell mechanics model (37) to describe process like growth, mitosis and motility. A more accurate description of the driving force, which accounts for the shape of the bleb and cell, requires a 2D model of blebbing. We present this in subsequent sections. In what follows, we describe the formulation of our time-dependent mechanical parameters, *k*_*b*_ (*t*), 𝜏_*c*_ (*t*) and 𝜏_*y*_ (*t*).

### Submodel for time dependent mechanical properties

In the local region where blebs form, the stiffness and viscosity of the boundary as well as the viscosity of the cytoplasm are very dynamic, as discussed previously. Initially, the separation of the membrane from the cortex creates a purely elastic boundary which expands under the force of pressurized fluid. As a new cortex reforms beneath the expanded membrane, the old cortex (actin scar) is simultaneously degraded. This alters the mechanical properties of the local bleb environment in two ways. First, the stiffness and viscosity of the boundary increases in proportion to the concentration of f-actin that has accumulated beneath the protruded membrane. Secondly, the degradation of the actin scar permits lager proteins, organelles and macromolecules to enter the cytoplasm between the protruded membrane and actin scar (bleb cytoplasm). This tends to increase the viscosity of the bleb cytoplasm in proportion to the concentration of f-actin remaining in the actin scar. Let *k*_*b*_ be the boundary stiffness due to the membrane and cortex, then we can model it as

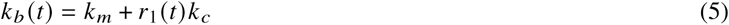

where 0 ≤ *r*_1_(*t*) ≤ 1 accounts for the proportion of cortex that has reformed beneath the protruded membrane. Likewise, the viscosity of the bleb boundary can be modeled as

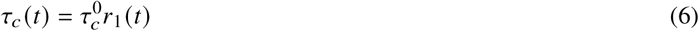

where 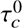 is the viscosity coefficients of the pre-bleb cortex. When the membrane first detaches from the cortex, the fluid in the bleb cytoplasm is less viscous due to the absence of larger proteins and organelles that could not pass through the cortex. Let 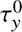 be the cytoplasmic viscosity prior to bleb expansion, then the initial viscosity of the bleb cytoplasm will be 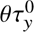 where < 𝜃 < 1. We expect the cytoplasmic viscosity to increase as more f-actin is removed from actin scar. Thus, we model the bleb cytoplasmic viscosity as

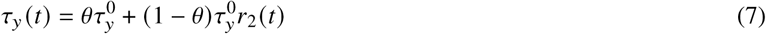

where *r*_2_(*t*) is the proportion of degraded actin. The precise form of the ratio *r*_1_(*t*) and *r*_2_(*t*) are determined using our recent data-driven modeling of cortex reformation (7). We set 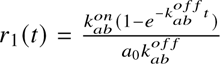 as the relative concentration of actin in the reformed cortex. Here, 𝑎_0_ is the actin concentration in the mature cortex, 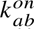 and 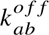 are the polymerization and depolymerization rate of actin in the reforming cortex. Similarly, 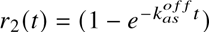 is the relative concentration of actin removed from the degrading cortex, with 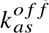 as the depolymerization rate of actin in the degrading cortex. Details on the derivation of *r*_1_(*t*), *r*_2_(*t*) are provided in the supplemental methods.

Our final model for the time dependent mechanical parameters for the cytoplasmic viscosity, boundary viscosity and boundary stiffness (due to membrane and cortex) thus become

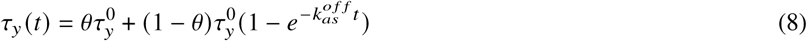

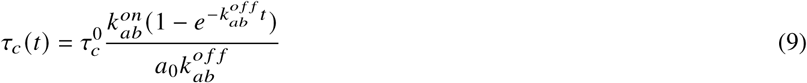

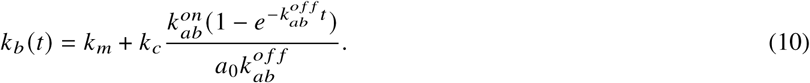

Taken together, Eqns. 3, 4,8, 9, 10, describe the 1D dynamics of bleb expansion. We note that whereas our model for radial displacement of the cell boundary in Eq. 3 is similar to the model for boundary displacement during micropipette aspiration in (38), it is distinct because it accounts for the resistance of linker proteins through the term *k*_𝑎_ 𝜌_𝑎_ and couples the viscous element of the cytoplasm in parallel with the viscoelastic element of the cell boundary. The combination of Eq. 3 and Eq. 4 to describe 1D bleb expansion is also unique to this work. Additionally, we have included time-dependent mechanical parameters (Eq. 8-Eq. 10) to capture dynamic changes in the structure of the cell boundary as a new cortex reforms and the old cortex is degraded. Additionally, whereas traditional models of bleb formation compute stretching forces between the cortex and membrane in order to detach linker proteins, our framework uses the displacement of the bleb boundary (𝑢) from its original location to compute the unbinding rate of linker proteins and hence the density of linker proteins 𝜌_𝑎_ available to resist further detachment with stiffness 𝜌_𝑎_ *k*_𝑎_.

### Numerical simulation of bleb expansion in 1D

We solve the 1D model using MATLAB’s ode23s solver. Given an applied pressure 𝐹_0_, we simulate bleb expansion by first determining the boundary displacement and linker density at which the cell boundary and cytoplasmic resistance would balance out the driving pressure. We refer to these as the critical displacement and critical density, denoted by 𝑢^0^ and 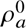 respectively and illustrated in Fig. 1C. These critical values are necessary to account for the cell’s initial attempt to re-equilibrate forces when the intracellular pressure is increased above resting levels. Note that when this initial displacement is insufficient to balance out the excess pressure, linker proteins are detached in a local region, the membrane separates from the cortex and a bleb is born (4). Thus, we initiate blebs by setting these critical values as the initial condition for our ode model (Eq. 3 - Eq. 4) and then mimic the detachment of linker proteins by setting 𝜌_𝑎_ = 0. The critical displacement and density are obtained by solving for the steady-states of Eq. 3 and Eq. 4 with constant mechanical parameters, since cortex reformation and degradation would not have begun at this time. Consequently, we obtain the following nonlinear solution

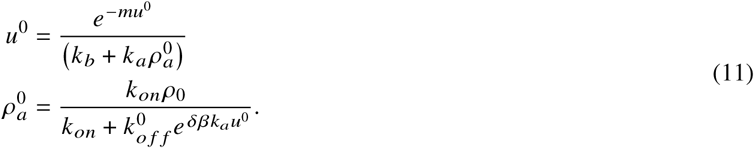

which can be solved using MATLAB’s inbuilt nonlinear equation solver. Details on how the steady-state values of the model were calculated has been provided in the supplemental methods. This idea of computing a critical displacement and linker density prior to membrane separation is new to this work and different from existing approaches for initiating blebs (27, 29, 39, 40).

### 2D dynamic model for bleb expansion

Whereas a 1D model of bleb expansion provides a system of ODE which are relatively easy to analyze and simulate, it does not provide a full description of the driving pressure that generates blebs or the dynamics of linker proteins as the membrane peels away from the cortex. Using the 1D model as a foundation, we build a model of blebbing in two-dimensions (2D). Inspired by the work of Iglesias (38), we describe the mechanical response of the cell to applied intracellular pressure as equivalent to an array of radial 1D viscoelastic elements, illustrated in Fig. 1B. We assume further that at each time *t*, the displacement vector 𝐼 (*x*, *y*, *t*) ∈ R^2^, varies continuously along the cell boundary, with magnitude in the normal direction given by the 1D displacement 𝑢(*t*). Here, *t* = *t*(*S*, *t*), *y* = *y*(*S*, *t*) are the coordinates of the cell boundary Γ(*t*) := {(*t*(*S*, *t*), *y*(*S*, *t*)) : *S* ∈ [𝑎, *b*]}. Similarly, we assume that the density of linker proteins, denoted by Φ(*x*, *y*, *t*), varies continuously along the cell boundary. In order to compute the gradient of these quantities, we need them to be defined in a neighborhood around the cell boundary. Consequently, we extend 𝐼 and Φ to be defined off the boundary by assuming that they are constant in the unit normal direction (𝑛^) to the interface. With this assumption, we obtain the following 2D model for bleb expansion

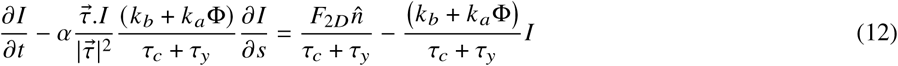

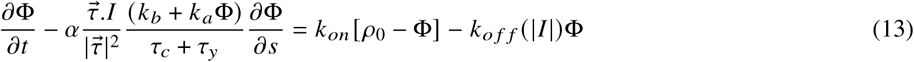

with spatially varying time dependent mechanical parameters,

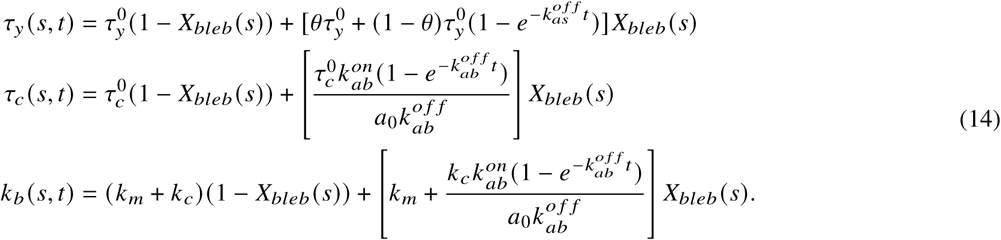

Here 𝜏® is the tangent vector to a point on the cell boundary, Γ. The spatiotemporal mechanical parameters 𝜏*c* (*S*, *t*), 𝜏_*y*_ (*S*, *t*) and *k*_*b*_ (*S*, *t*) are 2D extensions of the time dependent mechanical parameters described in Eq. 8-Eq. 10. The characteristic function 𝑋_*b*𝑙𝑒*b*_ (*S*) is 1 in the blebbing region and zero otherwise. Consequently, the mechanical parameter are only updated within the bleb, remaining constant outside the blebbing region. Note that Eq. 12 is similar to Eq. 3 with an additional term 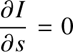 which accounts for spatial variation in the displacement of the boundary within the blebbing region, and vanishes wherever the displacement 𝐼 is uniform. Observe that in regions where 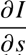, this term disappears and Eq. 12 simply becomes a vectorized form of Eq. 3. Likewise at points where 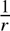 is large, mostly at the shoulder points of the bleb (transition point between detached bleb and attached cell boundary), this term increases the velocity of the boundary and consequently increases the likelihood of peeling (progressive widening of the bleb neck). A similar term appears in the linker density equation (𝐸𝑞. 13), playing a similar role. The dimensionless parameter 𝛼 ≥ 1 is a peeling factor which regulates the contribution of spatial gradients in boundary displacement and linker density to peeling of the bleb.

### Submodel for 2D driving pressure

One of the limitations of our 1D model is the pressure term, which is quite phenomenological. In 2D, we model the driving force 𝐹_2𝐷_ as

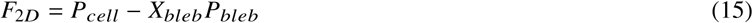

where 𝑃_*c*𝑒𝑙𝑙_ is the cytoplasmic pressure in the cell and 𝑃_*b*𝑙𝑒*b*_ is the pressure inside the bleb. We assume this pressure is constant within the cell (as was done in a recent cell mechanics model (37) to describe process like growth, mitosis and motility) and model it following Laplace’s Law which states that the pressure across a circular interface with curvature ^1^ subject to boundary tension 𝛾 is 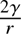. We extend this to non-circular interfaces as 2𝛾𝜅, where 𝜅 is the boundary curvature. Blebbing cells tend to preserve a constant cell volume(39). To enforce this, we introduce a volume pressure term, through the bulk modulus and cell volume, that decreases (increases) the cytoplasmic pressure if the cell volume increases (decreases). Additionally, we introduce a pressure term, *P_start_* to capture any excess intracellular pressure induced by say actomyosin contraction before bleb initiation. Thus, the cytoplasmic pressure *P_cell_* takes the form

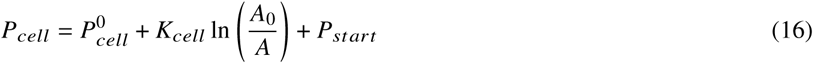

where, 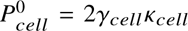 is the resting hydrostatic pressure of the cell with 𝛾_*c*𝑒𝑙𝑙_ and 𝜅_*c*𝑒𝑙𝑙_ as the boundary tension and curvature. The boundary tension 𝛾_*c*𝑒𝑙𝑙_ is taken as the sum of the membrane and cortical tensions, 𝛾*_m_* (*nN*/*μm*), 𝛾_*c*_ (*nN*/μ*m*) respectively. The bulk modulus of the cytoplasm is denoted as 𝐾_*c*𝑒𝑙𝑙_ (*nN*/μ*m*^2^) with 𝐴_0_, 𝐴 as the initial and current cell area. Observe that we can use the cell cross sectional area instead of volume in the volume constraint term because of our initial geometric assumption. We model the bleb pressure similarly as

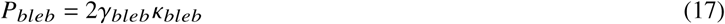

where 𝛾_*b*𝑙𝑒*b*_ (*nN*/μ*m*), 𝜅_*b*𝑙𝑒*b*_ represent the tension and curvature of the bleb boundary. Here, we set 𝛾_*b*𝑙𝑒*b*_ to be the membrane tension 𝛾_*m*_. Again, a detailed derivation of all models in Eq. 12 - Eq. 17 is provided in the supplemental methods.

### Numerical simulation of bleb expansion in 2D

The 2D model is solved by first discretizing all spatial derivatives using finite difference methods in order to convert it to a system of ordinary differential equations. The resulting system of ordinary differential equations is solved using MATLAB’s ode23s. Additional details on the numerical discretization of the 2D model have been provided in the supplementrary information. The process for simulating bleb expansion is illustrated in Fig. 1C. Prior to bleb initiation, we assume a resting intracellular pressure 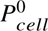 is balanced by the resistance of the viscoelastic boundary and linker proteins (with base line mechanical parameters). Upon the application of some excess pressure *P_start_* the cell boundary expands in an effort to equilibrate the applied pressure. This sets the displacement vector 𝐼 and linker density Φ to some critical value. To obtain these critical values, we assume there is initially no spatial variation in the boundary displacement 𝐼 and critical density Φ around the cell. Thus, 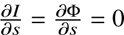 in the 2D model. Assuming further that any initial boundary displacement occurs in the normal direction (𝑛^), we can set 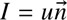, which allows us to use the 1D critical displacement 𝑢^0^ and linker density 𝜌^0^ as initial conditions for the 2D model. Linker proteins are subsequently detached from a set region of the cell boundary to observe bleb expansion. The practice of presetting the local region where blebs form is common in recent models of bleb expansion (27, 28).

## EXPERIMENTAL RESULTS

We investigated the role of TalA in regulating blebbing by analyzing bleb characteristics (frequency and area) from wild type *Dictyostelium discoideum* cells (Ax2) and *talA* null cells, chemotaxing under an agarose block towards Cyclic-AMP (cAMP). We use this experimental set up because cAMP is the chemoattractant that Dictyostelium cells respond to during aggregation of individual cells to form a multicellular fruiting body. Blebs have been implicated as the dominant mode of migration during this period, when cells transition from migrating in two-dimensions to a three-dimensional environment (6). Additionally, the agarose block offers a confined environment that mimics movement in a three-dimensional environment to help induce blebbing. See Fig. 2A for an illustration of the under-agarose setup used for experiments and Materials and Methods for a complete description of the assay.

**Figure 2:**
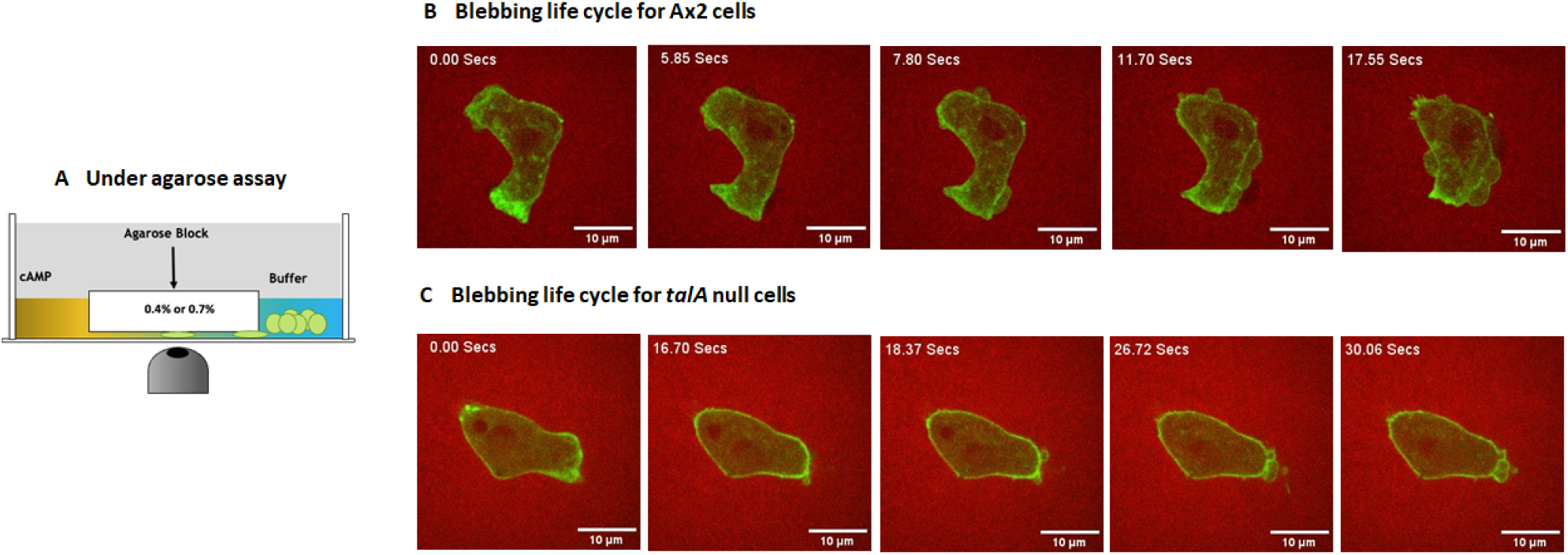
Blebbing Ax2 and *talA* null cells. A) Illustration of underagarose assay used to collect data on blebbing chemotaxing cells. An illustration of b) wild type cell and c) *talA* null cell blebbing under 0.7% agarose. Factin is marked by LifeAct-GFP and the membrane is shown as a shadow against the agarose gel lased with RITC-dextran (red)

Cells were observed under low confinement (0.4% agarose) and high confinement (0.7% agarose) for a duration of 30 seconds each. An example of blebbing in Ax2 and *talA* null cells is shown in Fig. 2(B,C) as well as in the supplemental movies. Notice the complete bleb cycle in Ax2 cells of membrane separation (5.85 secs), cortex reformation and accumulation of F-actin inside the bleb (7.80 secs) and complete degradation of the cortex at 17.55 secs, which is consistent with previous published work. A similar cycle can be observed in *talA* null cells. The total number of cells analyzed and the number of blebs produced by these cells are provided in Table 1. The data represents observations pooled from two independent experimental trials. Bleb characteristics were measured after the cortex, visualized with LifeAct-GFP, had fully formed around the protruded membrane in both Ax2 and *talA* null cells.

**Table 1:**
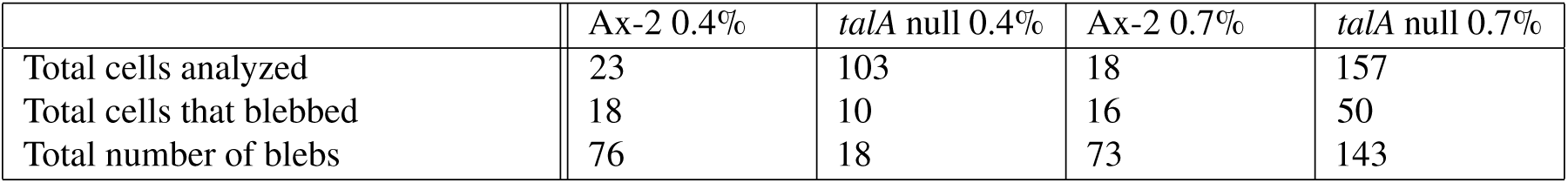
Summary of experimental data. This table summarizes the cellular data analyzed. The “Total cells analyzed” reflects counts for all cells (in frame and not in contact with other cells) that were used. The “Total cells that blebbed” is the number of cells that had at least one bleb.

|  | Ax-2 0.4% | *talA* null 0.4% | Ax-2 0.7% | *talA* null 0.7% |
| --- | --- | --- | --- | --- |
| Total cells analyzed | 23 | 103 | 18 | 157 |
| Total cells that blebbed | 18 | 10 | 16 | 50 |
| Total number of blebs | 76 | 18 | 73 | 143 |

### *talA* null cells produce fewer blebs than wild type

We analyzed the proportion of wild type (Ax2) and *talA* null cells that produced blebs under low and high confinement. A total of 23 Ax2 cells were analyzed under low confinement conditions with 78% of them producing at least one bleb (Table 1). On the contrary, just under 10% of the *talA* null cells bleb. A similar pattern can be observed under high confinement. Whereas about 90% of Ax2 cells produce at least one bleb, just about 30% of *talA* null cells bleb. We compared the number of blebs produced per cell in Ax2 and *talA* null cells. As illustrated in Fig. 3A, *talA* null cells produce fewer blebs than Ax2 under all confinement conditions. Whereas Ax2 cells produce a median of 4 blebs per cell, under low confinement, the median bleb frequency in *talA* null cells is zero under similar conditions. Both cells appear to increase their bleb frequency in response to confinement. Contrary to expected results, these data suggest that *talA* null cells bleb less frequently than wild type under any confinement condition, with a higher proportion of them blebbing in environments with higher confinement.

**Figure 3:**
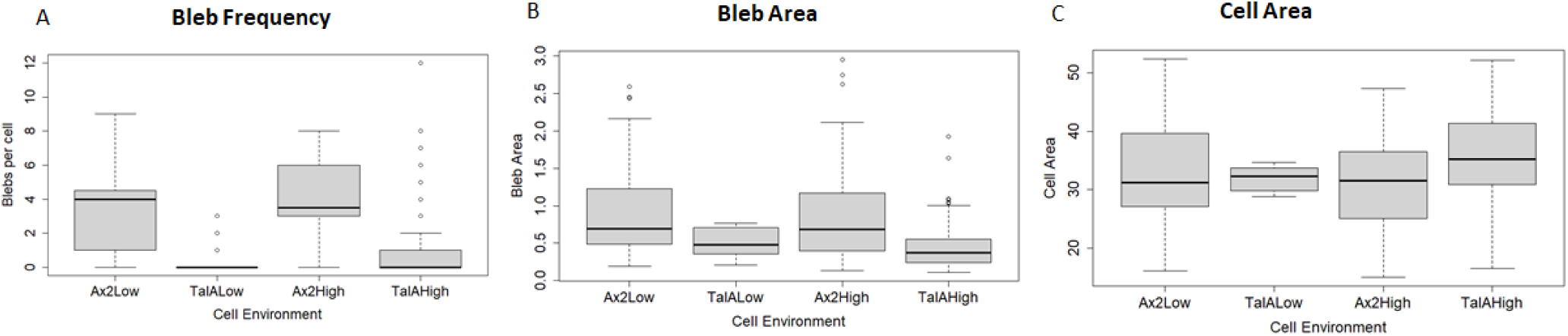
Experimental data. Data on a) frequency of blebs per cell b) bleb area c) Cell area for both wild type (Ax2) and mutant (*talA* null) cells shown in Table 1. Ax2High ( wild type under high confinement), Ax2Low (wild type under low confinement), TalAHigh (*talA* null under high confinement) and TalALow (*talA* null under low confinement).

### *talA* null cells have smaller blebs and larger cell surface area than wild type

Blebs are driven by intracellular pressure which is regulated by actomyosin contractility of the cortex. Tinevez et al. have identified the existence of a critical cortical tension (i.e intracellular pressure) above which cells form blebs (18). Since *talA* null cells rarely bleb, we reasoned that this could result from a low intracellular pressure. Bleb size correlates with intracellular pressure (5, 28), therefore it can act as a gauge for the amount of pressure in the cell (41). Thus, we tested our hypothesis by measuring the size of blebs produced by *talA* null cells and compared it to bleb size of Ax2 cells. Bleb size is characterized as the cross sectional area of the visible bleb from our microscopy images. See Materials and Methods for details on how area measurements were taken.

The distribution of bleb area measurements is shown in Fig. 3B. Notice that, *talA* null cells have smaller bleb area than wild type under all confinement conditions. The decrease in bleb size under high confinement was found to be statistically significant (pvalue <0.01) whereas that for low confinement was not (pvalue=0.08). Bleb size remains fairly constant for each cell line under different levels of confinement (p-value=0.63: Ax2, p-value=0.17: *talA* null). These findings suggest that *talA* null cells have lower intracellular pressure, with a greater reduction in pressure observed in environments with high confinement.

Next we ask how TalA could regulate intracellular pressure. Fluid pressure in the cell is inversely proportional to the surface area to volume ratio of the cell. *Dictyostelium* cells are known to maintain an almost constant fluid volume during migration (42), therefore changes in the surface area of the cell could reduce the intracellular pressure. We sought to determine whether *talA* null cells have increased surface area, which would explain the low intracellular pressure. To achieve this, we compared cross sectional area (2 dimensional) of *talA* null cells to Ax2 under different confinement conditions. Our results, presented in (Fig. 3C) show that *talA* null cells have a larger cross sectional area than Ax2 under high confinement (p-value<0.01). No significant difference in cell area was observed under low confinement. This data supports our hypothesis that the reduced pressure in *talA* null cells is driven by an increase in the surface area of the cell.

## THEORETICAL RESULTS

The question that remains is why a loss of TalA would result in an increase in the surface area of the cell. We reasoned that knocking out *talA* leaves the membrane weakly bound to the cortex by TalB. This weaker membrane-cortex attachment results in a uniform extension of the membrane away from the cortex when the cell is pressurized, creating an initial pressure relief that limits the prevailing pressure for blebbing. In recent work by Tsujioka et al.(20) *Dictyostelium* cells devoid of TalA left membrane tails in the cell posterior, supporting the idea that the membrane is not as tightly bound to the cortex when TalA is absent. We have developed mathematical models of bleb expansion to directly test the hypothesis that decreasing the strength of linker proteins lowers the prevailing pressure for bleb expansion by uniformly pushing the membrane from the cortex. We also test that this decrease in pressure is sufficient to produce smaller blebs, as observed in the experimental data. The models are developed in the Materials and Methods section, with additional details provided in the supplemental methods.

The first model in Eq. 3 – Eq. 4 tracks bleb expansion in one-dimension (1D) along a radial line drawn through the bleb using a phenomenological pressure term (Fig. 1A, bottom). This simple 1D framework allows for an analysis of the factors influencing the steady-state bleb size as well as the pre-bleb mechanical state of the cell, upon the application of excess pressure. The second model in Eq. 12 – Eq. 14 is a two-dimensional model developed to track the expansion of realistic bleb shapes using a more biologically relevant expression for intracellular pressure.This is the primary model for testing our hypothesis since the 2D framework permits an investigation of the effect of weakening linker proteins on the surface area of the cell. We did not attempt to obtain a steady-state model of bleb size in 2D, as was done in 1D, since this is non-trivial given the complexity of our bleb expansion model. Rather, we simulate the 2D model up to a time when bleb expansion is known to stabilize experimentally. A necessary part of our model’s complexity comes from our efforts to replicate realistic bleb expansion by accounting for the reformation and degradation of the cortex.

First, we validate our mathematical models by exploring their ability to replicate known effects of cell mechanical parameters on bleb size and then apply them to test our stated hypothesis. We also use these models to investigate how actin dynamics regulates bleb size, since cortex reformation and degradation are observed during bleb stabilization. Recent experimental data on the dynamics of bleb formation in *Dictyostelium discoideum* cells suggests that blebs reach maximum size within the first 5-10 seconds after membrane separation and stabilize by 25 seconds (7). Therefore, we simulate our models (first 1D and then 2D) for 10-25 seconds. All model parameters used in the simulation are presented in Table 2, with their corresponding units and references. Since the 1D model is the foundation for our 2D model, we validate that first before validating the full 2D model. MATLAB codes for generating all the simulated results are available on GitHub via the link https://github.com/easantea/BlebExpansion.git.

**Table 2:** Parameter values for simulating the bleb expansion models.

| Parameter | Description | Value | References |
| --- | --- | --- | --- |
| $F_0$ | Resting Intracellular pressure | $0.08 \text{ nN}/(\mu\text{m})^2$ | (39) |
| $m$ | Rate at which driving pressure decreases in 1D model | 0.1 | This work |
| $r$ | Radius of the cell | $5 \mu\text{m}$ | This work |
| $\tau_c^0$ | Viscosity coefficient mature cortex | $0.064 \text{ nNs}/(\mu\text{m})^3$ | (38) |
| $a_0$ | Actin density in mature cortex | 1.8 | (7) |
| $k_{ab}^{on}$ | Actin polymerization rate at bleb boundary | 0.7602 | (7) |
| $k_{ab}^{off}$ | Actin depolymerization rate at bleb boundary | 0.4344 | (7) |
| $k_{as}^{off}$ | Actin depolymerization rate at actin scar | 0.0120 | (7) |
| $\tau_y^0$ | Viscosity coefficient of cytoplasm | $6.09 \text{ nNs}/(\mu\text{m})^3$ | (38) |
| $K_{cell}$ | Bulk modulus of the cell body | $0.225 \text{ nN}/(\mu\text{m})^2$ | (39) |
| $k_c$ | Spring constant for the cortex | $0.098 \text{ nN}/(\mu\text{m})^3$ | (38) |
| $k_m$ | Spring constant for membrane | $0.00392 \text{ nN}/(\mu\text{m})^3$ | Estimate using (38) |
| $\gamma_m$ | Membrane tension | $0.032 \text{ nN}/(\mu\text{m})$ | Estimate using (28) |
| $\gamma_c$ | Cortical tension | $0.2 \text{ nN}/(\mu\text{m})$ | Estimate using (28) |
| $\theta$ | Ratio of membrane stiffness to cortical stiffness | 1/250 | (28) |
| $k_a$ | Strength of single linker protein | $10^{-1} \text{ nN}/(\mu\text{m})$ | (35) |
| $\rho_0$ | Total available density of linker protein | $10^2 (\mu\text{m})^{-2}$ | (35) |
| $k_{on}$ | Binding rate of linker protein | $10^4 (\text{s})^{-1}$ | (35) |
| $k_{off}^0$ | Detachment rate of linker proteins at zero displacement | $10 (\text{s})^{-1}$ | (35) |
| $\beta$ | $1/\text{Thermal energy } (k_B T)^{-1}$, $k_B$ is Boltzmann constant and $T$ is thermodynamics temperature | $\frac{10^6}{4.1} ((\text{nN})(\mu\text{m}))^{-1}$ | (35) |
| $\delta$ | characteristic bond length | $10^{-3}(\mu\text{m})$ | (35) |
| $\alpha$ | Peeling factor | [1, 6] | This work |

### Effect of cell viscoelastic properties on bleb size

When the membrane first detaches from the cortex it expands under the force of flowing cytosol, driven by intracellular pressure and resisted only by its stiffness, *k*_*m*_and initial fluid viscosity 𝜏_*y*_. As the cortex begins to reform beneath the membrane and degrade at the old location of the cortex, the boundary resistance increases and slows down bleb expansion. It is well known that bleb size increases with increasing intracellular pressure (or environmental confinement) and decreases as the viscoelastic properties of the cortex and membrane increases (5, 6, 28). It has also being observed experimentally that as actin reforms beneath the protruded membrane, the bleb boundary undergoes a partial retraction as it approaches steady-state size (7). We sought to test our model’s ability to capture these known trends by varying our model parameters. The size of blebs in the 1D model is taken to be the displacement 𝑢 at a fixed point of the cell boundary, along a radial line drawn through the boundary. We allowed the cortex-free bleb to expand for 2 seconds before beginning cortex reformation, as was determined from experimental data in our previous study (7).

Our results in Fig. 4 and Fig. 5 both demonstrate the expected dynamics of a bleb. That is, a rapid expansion within the first 2 seconds and slower expansion or retraction to an equilibrium size after cortex reformation is initiated. We varied the resting intracellular pressure 𝐹_0_ and membrane stiffness *k*_*m*_, cortex stiffness *k*_*c*_ and baseline cortex viscosity 𝜏^0^ by scaling them from to 10 times their basal values given in Table 2. Observe from Fig. 4A that bleb size increases with intracellular pressure. Similarly, as membrane elastic stiffness decreases, the bleb expands faster to a higher maximum size and then retracts to a larger equilibrium size (Fig. 4B). As the cortex stiffness decreases, blebs expand faster and settle down to a larger equilibrium size (Fig. 4C). Interestingly, there appears to be two separate bleb expansion regimes regulated by cortical stiffness, one in which blebs partially retract to equilibrium size, after reaching their maximum size, and the other where blebs do not retract, but expand gradually to their equilibrium size. Surprisingly, varying cortex viscosity, does not alter equilibrium bleb size, rather it appears to regulate the speed of expansion with a less viscous cortex leading to a larger maximal bleb size (Fig. 4D). Together, these results support the known behavior of expanding blebs and serve as a semi-quantitative validation of our 1D model.

**Figure 4:**
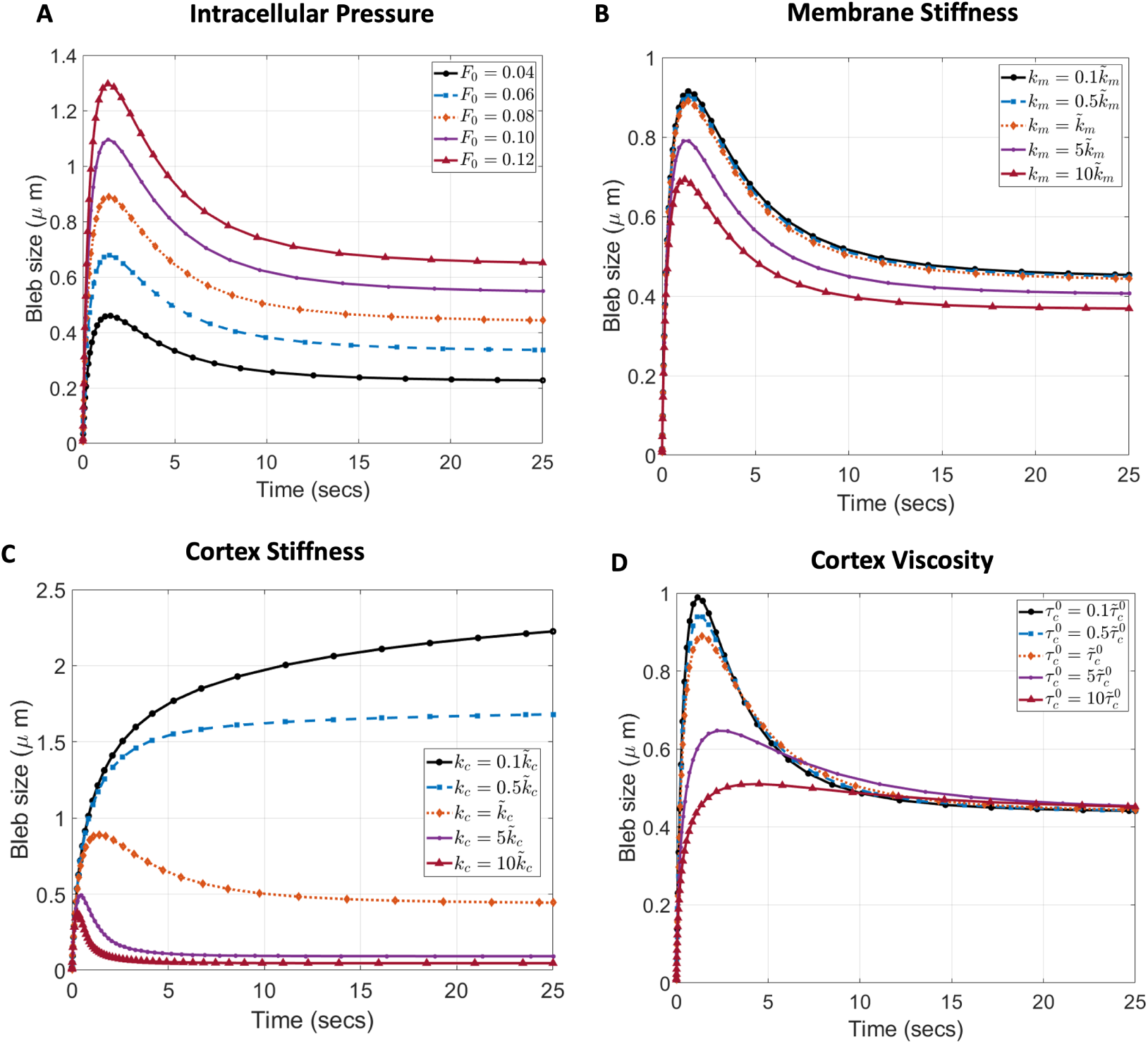
Influence of cell mechanical properties on bleb size. Effect of A) Intracellular pressure B) membrane stiffness, C) cortex stiffness and D) cortex viscosity. Bleb size is measured as the strain of the cell boundary at a single point. ̃*r* is the baseline value of parameter *r* reported in Table 2.

**Figure 5:**
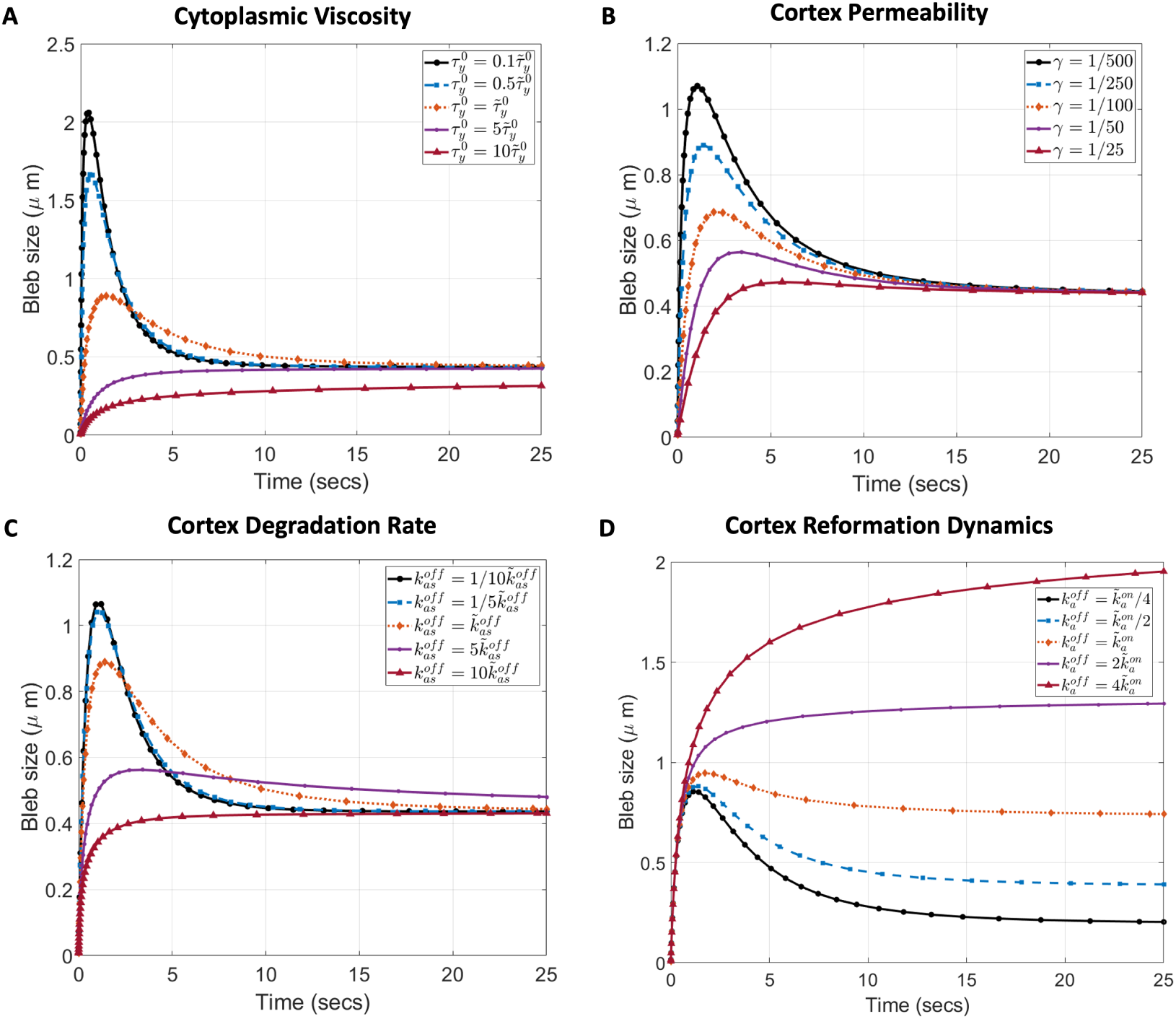
Influence of viscoelastic properties and actin dynamics on bleb size. Effect of A) cytoplasmic viscosity B) cortex permeability, modeled as the fraction of actin concentration in the cortex (actin scar) at the onset of blebbing C) degradation rate of actin scar (old cortex), D) ratio of actin polymerization to depolymerization rate at the reforming cortex (bleb cortex), on bleb size. Bleb size is measured as the strain of the cell boundary at a single point. ̃*r* is the baseline value of parameter *r* reported in Table 2.

### Effect of actin dynamics, cortex permeability and cytoplasmic viscosity on bleb size

We found it interesting that the equilibrium bleb size did not depend on the cortex viscosity but rather on membrane and cortex stiffness as well as intracellular pressure. Curious about whether this is supported theoretically and also to identify other model parameters influencing the equilibrium bleb size, we analyzed the equilibrium solutions of the 1D model (Eq. 3) with 𝜌_𝑎_ = 0 and time dependent parameters described in Eq. 8-Eq. 10. Note that by setting 𝜌_𝑎_ = 0, we remove linker proteins from equilibrium analysis since they are detached when the membrane separates from the cortex. In order to perform equilibrium analysis, we converted the 1D model (Eq. 3), with 𝜌_𝑎_ = 0, to an autonomous system of differential equation by writing separate rate equations for all time dependent parameters described in Eq. 8-Eq. 10. The resulting steady-state bleb size 𝑢^∗^ satisfies the condition

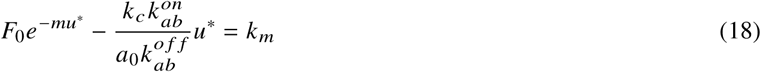

which confirms the dependence of bleb size (𝑢^∗^) on initial intracellular pressure, 𝐹_0_, cortex stiffness *k*_*c*_ and membrane stiffness *k*_*m*_, observed from our model simulations. Additionally, the condition (Eq. 18) points to a dependence of equilibrium bleb size on actin reformation dynamics through the ratio of actin polymerization and depolymerization rate 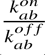. However, the degradation rate of the actin scar (old cortex), cytoplasmic viscosity and cortex permeability have no effect on the equilibrium bleb size. Details of the autonomous reformulation of the model and derivation of the condition Eq. 18 for equilibrium bleb size can be found in the supplemental methods.

Next, we wanted to understand how cortex reformation affects the final bleb size and then explore the effect, if any, that cortex degradation, cytoplasmic viscosity and cortex permeability may have on bleb expansion dynamics. Our results, shown in Fig. 5, suggest that a more viscous cytosol will cause blebs to expand monotonically to their resting size, without retracting (Fig. 5A). As the viscosity of the fluid reduces, blebs have a greater initial expansion before retracting to their equilibrium size. Cortex permeability and degradation rate of the actin scar have a similar effect on bleb expansion dynamics (Fig. 5B,C). As predicted theoretically, none of these factors alter the equilibrium size of blebs. In support of our equilibrium analysis, we see that varying the dynamics of cortex reformation changes the equilibrium bleb size (Fig. 5D). Interestingly, partial bleb retraction to equilibrium size seems to be regulated by the relative rate of actin polymerization 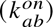 and depolymerization 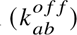 at the bleb cortex. Particularly, when 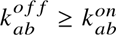 no retraction occurs and the blebs expand monotonically to an equilibrium size. On the contrary, blebs retract before reaching equilibrium size when 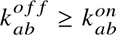.

### Influence of linker protein strength on bleb size

Next, we test whether decreasing linker protein strength can decrease bleb size. To this end, we analyzed how linker protein stiffness *k*_𝑎_ affects the critical displacement of the cell boundary and subsequently bleb size. Recall that the critical displacement is the maximal displacement of the cell boundary prior to detachment of linker proteins, in response to an increase in intracellular pressure. This critical displacement sets the initial condition for our 1D model, and thus determines the pressure available for bleb expansion. We obtain this value by finding the equilibrium solution of our 1D model (Eq. 3-Eq. 4) with constant viscoelastic parameters, as shown in Eq. 11. Since the uniform expansion of the cell membrane (around the entire cell) does not constitute a bleb, we calculate bleb size by subtracting the boundary displacement after 25 seconds (when the bleb has stabilized) from the corresponding critical displacement (before membrane separation).

Fig. 6 shows the bleb formation dynamics under high and low pressure for linker stiffness ranging from (0.005 − 0.2) *nN*/μ*m*. Notice that the equilibrium bleb size decreases as the linker protein stiffness decreases with larger blebs forming when pressure is higher (Fig. 6A,B). These results agree with our experimental observations in Fig. 3B where smaller blebs form in the absence of TalA proteins. Note from Fig. 6C that the critical displacement increases as the linker stiffness decreases. Our phenomenological model for intracellular pressure available for bleb expansion is 𝐹_1𝐷_ = 𝐹_0_𝑒^−*m*𝑢^ which decreases as the critical displacement increases. Thus, the smaller blebs from our model simulation can be explained by a decrease in the driving pressure, prior to bleb formation. Taken together, our 1D model supports our stated hypothesis by suggesting that weaker linker proteins result in a larger critical displacement which reduces the intracellular pressure required to form blebs. Note that, this conclusion could not have been obtained from our steady-state model in Eq. 18, which does not depend on the initial state of the system. Thus, our effort to explain the experimental data requires the solution of the full dynamic model.

**Figure 6:**
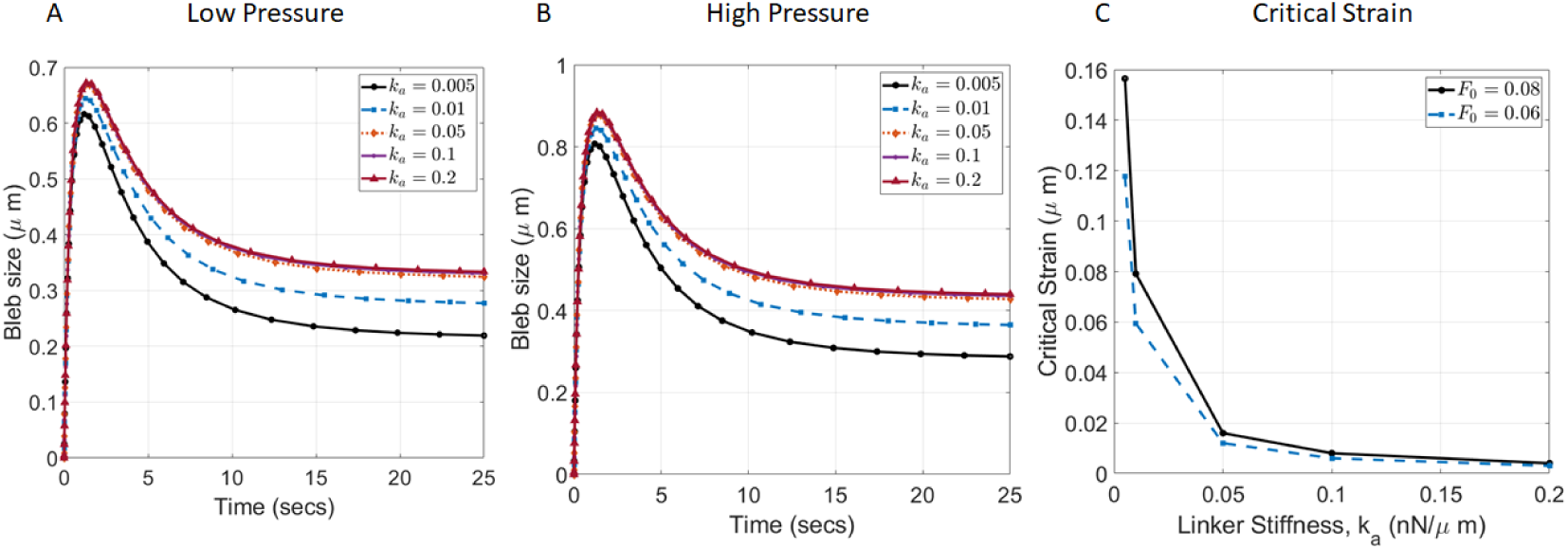
Influence of linker proteins strength on bleb size. Effect of linker protein strength of bleb size under A) low and B) high intracellular pressure. C) Critical displacement of cell boundary for different intracellular pressure. Curves are generated from 1D model.

Whereas our 1D model supports our stated hypothesis, it is based on a phenomenological model for intracellular pressure which does not take the cell shape into account. Our measurement of bleb size is also based on a radial displacement instead of the area of the bleb formed, as used in the experimental data. Thus, it is unclear if the same conclusions will be arrived at in a more geometrically relevant 2D setting. We explore this in the following sections.

### Cell area increases with decreasing linker protein strength

Our 2D model for bleb expansion is presented in Eq. 12– Eq. 14 in the Materials and Methods section along with details on how we generate blebs. We validated the model by testing known viscoelastic effects on bleb size. As blebs expand, the bleb neck (region of cell boundary where membrane detached from cortex) tends to increase. We also examined our model’s ability to generate this known phenomena.The bleb expansion dynamics from our 2D model over the first 10 seconds, can be observed in Fig. 7A. We simulated bleb expansion for 10 seconds instead of 25 seconds (done in 1D case) since our prior experiments (7) and 1D simulations (in this work) suggest that blebs achieve close to 90% of the equilibrium size within 10 seconds. This also helped to reduce the total run time for computational experiments without altering the results significantly.

**Figure 7:**
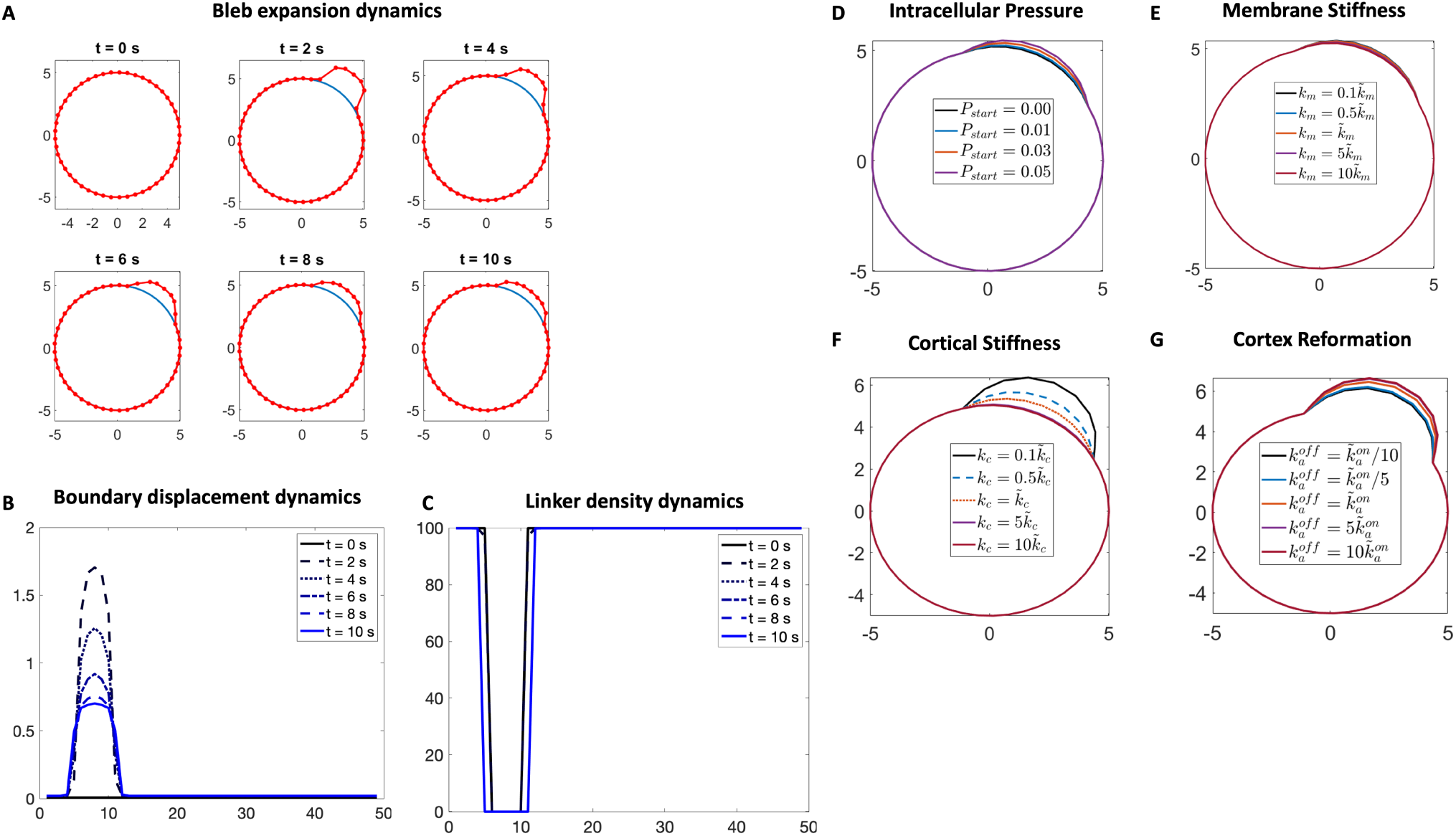
Validation of 2D blebbing model. A) Dynamics of bleb expansion for the first 10 seconds after membrane detachment with *P_start_* = 0.14 *nN*/μ*m*^2^ and peeling factor 𝛼 = 6. B) Boundary displacement and C) Linker proteins density of bleb in part A showing a widening bleb neck. Effect of D) intracellular pressure E) membrane stiffness F) cortical stiffness and G) cortex reformation on bleb size. ̃*r* is the baseline value reported in Table 2 of a parameter *r*. Peeling factor for D-G is set to 1.

Over the first two seconds, the membrane expands to a maximum bleb area and then slowly retracts to an equilibrium size, once the cortex begins to reform beneath the protruded membrane.The bleb neck can also be seen to widen in Fig. 7B,C as the bleb expands. The progressive widening of the bleb neck was only observed for peeling factor of 𝛼 = 6 and *P_start_* > 0.12*nN*/μ*m*^2^. For 𝛼 < 6 or 𝛼 = 6 and 0.004 < *P_start_* < 0.12*nN*/μ*m*^2^, the bleb only expands within the preset region used for initiation (Fig. S1). Interestingly, blebs fully retracted for *P_start_* < 0.004, indicative of the existence of some minimum pressure to form stable blebs.

Consistent with our 1D results, increasing the excess pressure *P_start_* leads to formation of larger blebs (Fig. 7D). Similarly, increasing membrane stiffness and cortical stiffness both reduce bleb size, with cortical stiffness having a greater effect (Fig. 7 E,F). In agreement with the 1D model, the dynamics of cortex reformation, in particular the ratio of actin polymerization to depolymerization rate, also affects bleb size (Fig. 7G).

We first examined the effect of linker protein strength on the critical area of the cell. This is the cell area when the prevailing intracellular pressure is balanced by the viscoelastic boundary, just prior to the detachment of linker proteins. Observe from Fig. 8A that the critical area increases as the linker stiffness is reduced, with a greater critical area for higher intracellular pressure. The difference in critical area between high and low pressure is smaller for stiffer linker proteins (*k*_𝑎_ ≥ 0.1) and increases as the linker stiffness is reduced (*k*_𝑎_ < 0.1). This observation agrees with our experimental results, where we had very little difference between the cell areas of Ax2 *Dictyostelium* cells under high and low agarose but a greater difference between *talA* null cells under similar conditions (Fig. 3C). Note that the range of simulated critical cell area is consistent with the area measured in our experimental data. In particular, the median cell area for Ax2 cells is around 32 μ*m*^2^ whereas that for *talA* null is around 35 μ*m*^2^ (𝐹𝑖𝑔. 3𝐶, high pressure).

**Figure 8:**
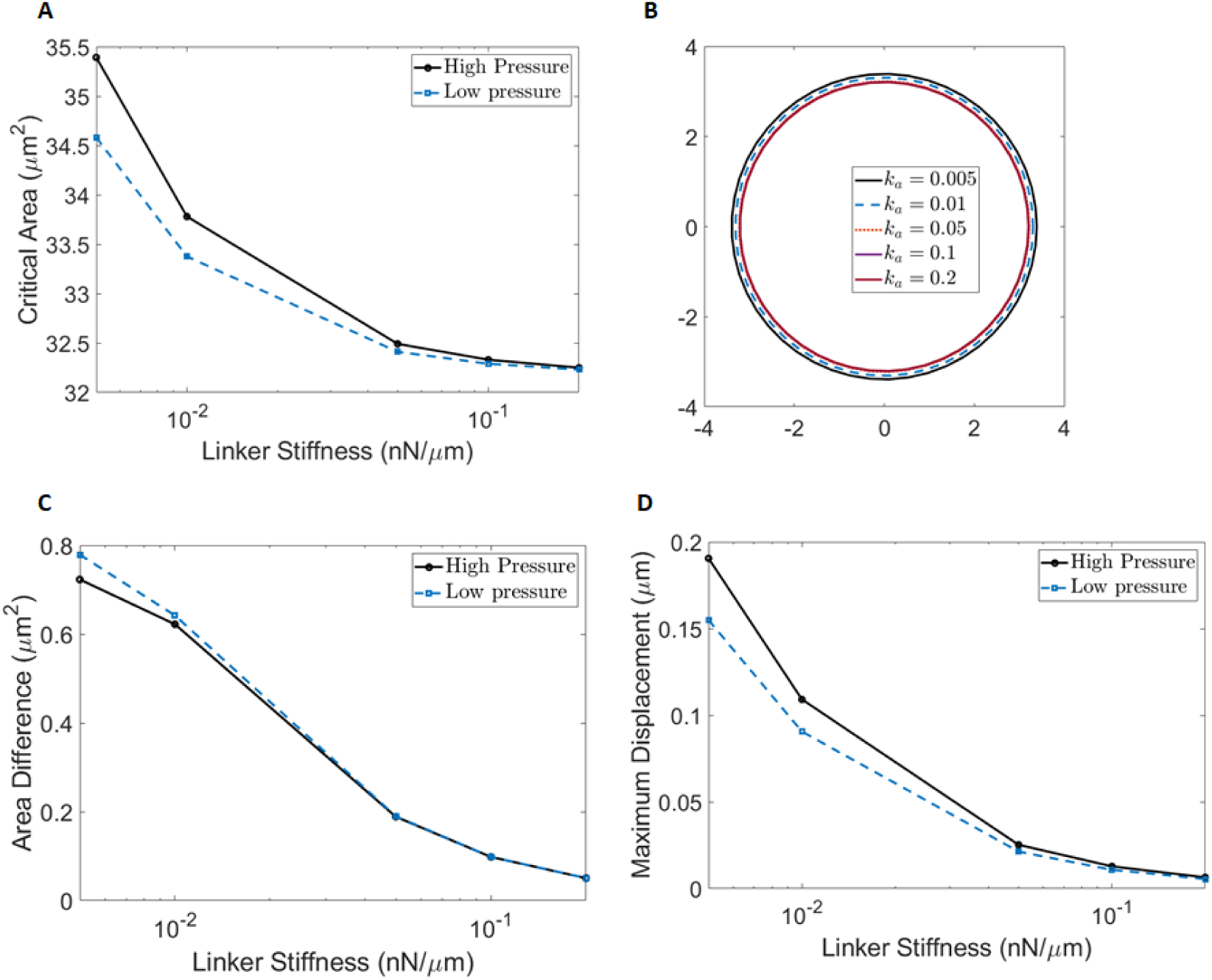
Response of cell boundary to excess pressure. Response of cell boundary to excess pressure of 0.05*nN*/(μ*m*^2^)), without detaching linker proteins, under varying linker protein strength over a 10 second duration. No blebs are initiated. A) Critical cell area prior to application of excess pressure. B) Final location of cell boundary after application of excess pressure when resting pressure is 0.08*nN*/(μ*m*^2^)). C) Difference in final area of cell and the critical area under high (0.08*nN*/(μ*m*^2^))) and low (0.06*nN*/(μ*m*^2^))) resting intracellular pressure. D) Maximal displacement of the cell boundary from its location at critical strain. Peeling factor 𝛼 = 1.

We also examined the effect that an excess pressure will have on the cell area and boundary displacement, without initiating blebs. This is important since our working hypothesis suggests that cell area increases resulting from reduced linker stiffness limits the driving pressure necessary for bleb expansion. To test this, we run our model for the same time duration (10 seconds), without detaching linker proteins (i.e initiating blebs), for different linker stiffness. Our results, shown in Fig. 8B, suggests that under the same intracellular pressure, the cell boundary is pushed out further in response to excess pressure when weaker linker proteins are present. Additionally, the maximum displacement of the cell boundary from its initial configuration is greater for high resting pressure than for lower pressure (Fig. 8D). It is note worthy that the difference between the final cell area (after application of excess pressure and no bleb formation) and the critical cell area (before application of excess pressure) is small relative to the scale of the critical area of the cell (Fig. 8C). Thus, our model does not yield a significant change in cell area in response to excess pressure, when blebs are not initiated.

### Bleb area decreases with decreasing linker protein strength

Next, we allowed blebs to form (by removing linker proteins from a local region of the cell boundary) and compared the final boundary profile (with a bleb) to the final profile without bleb formation (discussed in the previous section) as well as the critical profile (measured prior to the application of excess pressure). Our results in Fig. 9A qualitatively show that smaller blebs form as the stiffness of the linker protein is reduced. Notice that the cell boundary when blebs are not initiated in response to excess pressure (no-bleb boundary, dashed blue line in Fig. 9A) appears to overlay with the boundary when blebs form (bleb boundary, dotted orange line in Fig. 9A) everywhere except at the blebbing region. These two boundaries (bleb and no bleb) also appear to differ more from the critical boundary (solid black line in Fig. 9A) as the linker protein stiffness is decreased. Our simulations thus show that the cell boundary is uniformly pushed out in response to excess pressure everywhere (as shown in Fig. 8b), with a greater extension where linker proteins are detached (yielding blebs). Based, on this observation, we quantified the bleb area formed by subtracting the cell area enclosed by the bleb boundary (dotted orange line in Fig. 9A) from the area enclosed by the no-bleb boundary (dashed blue line in Fig. 9A). We can observe quantitatively that bleb area decreases as linker protein stiffness decreases, with larger blebs formed for higher pressure ( Fig. 9B). Notice that the size of blebs at our baseline value for linker stiffness (0.1nN/μ*m*) agrees with the median bleb area for wild type cells shown in our experimental data.

**Figure 9:**
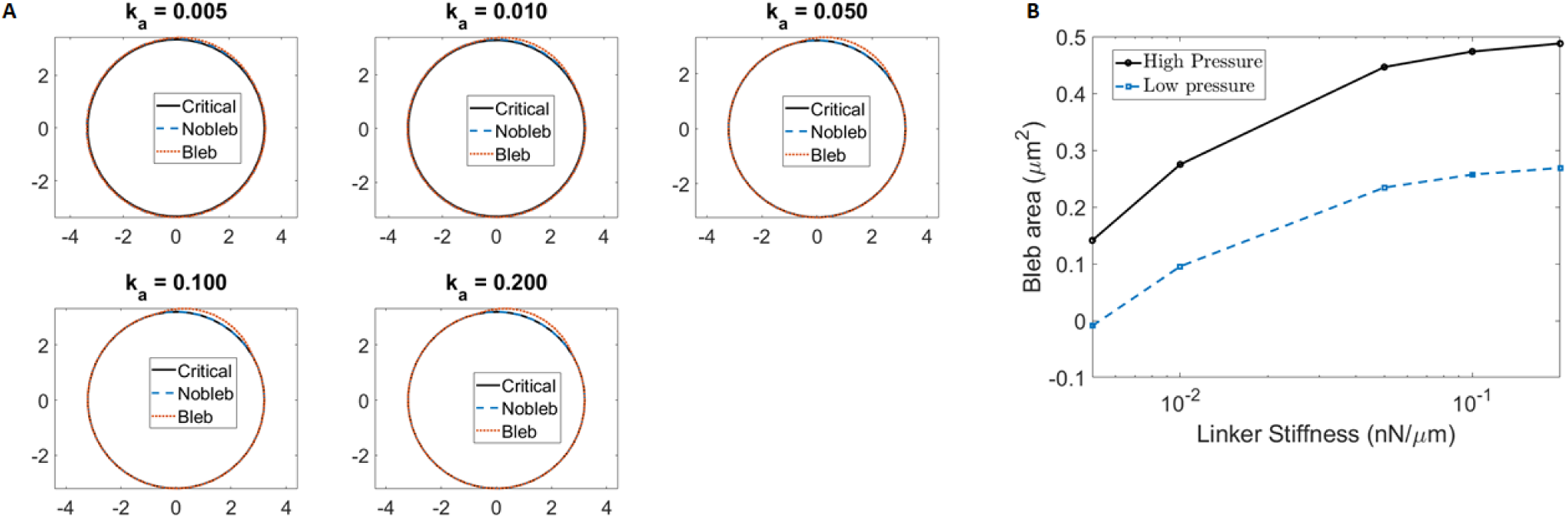
Effect of linker protein strength on bleb size. A) Bleb shape at different linker stiffness at high intracellular pressure. ‘Critical’ boundary (solid black line) is the critical cell boundary which balances the resting intracellular pressure. The ‘Nobleb’ boundary (dashed blue line) is the displaced boundary after application of excess pressure, without detaching linker proteins locally. The ‘Bleb’ boundary (dotted orange line) is the final cell boundary when blebs form. B) Bleb area for high and low resting pressure with varying linker stiffness. Peeling factor 𝛼 = 1.

### Effect of linker protein strength on bleb size under various viscoelastic conditions and actin dynamics

Finally, we wanted to examine how different elastic and viscous properties of the cell as well as actin dynamics affect the extent to which linker protein strength alters bleb size. This is an attempt to use our mathematical model to investigate how different cells (with different mechanical properties) might respond to weakening the stiffness of linker proteins. The mechanical properties examined are cortical stiffness, cortex viscosity, cortex permeability and cytoplasmic viscosity. We also investigated the influence of actin dynamics during cortex reformation and degradation. Each of these properties were varied over a high or low normal parameter range. High and low properties correspond to twice and half the baseline values of parameters presented in Table 2.

As can be observed from Fig. 10A,C,D,F cortical stiffness, cytoplasmic viscosity, cortex reformation rate and cortex degradation rate all influence the effect of linker protein strength of bleb size. At normal levels of of these parameters, bleb size is decreased by ∼ 70% over the range of linker protein stiffness examined. For cells with higher than normal cortical stiffness, blebs are generally smaller whereas they tend to be larger when the cortical stiffness is reduced below normal levels (Fig. 10A). Decreasing the cortical stiffness reduces the percentage loss in in bleb size to 36%, whereas blebs are completely lost for cells with higher cortical stiffness. The rate at which the cortex reforms during bleb stabilization also affects the overall impact of linker protein strength on bleb size. Generally we observe from Fig. 10C that cells with a faster cortex reformation rate tend to have smaller blebs.Slowing down cortex reformation reduces the percentage change in bleb size to 35% whereas blebs are completely lost when the reformation rate is doubled (made faster). For cytoplasmic viscosity, we observe from Fig. 10D a general increase in bleb size for cells with higher cytoplasmic viscosity. The percentage change in bleb size reduces to 42% for high cytoplasmic viscosity and increases to 83% for cells with lower viscosity. Increasing the speed at which the cortex degrades tends to increase bleb size (Fig. 10F). Increasing the degradation rate reduces the percentage change in bleb size to 56% whereas slowing down degradation does not significantly alter the percentage change compared to normal rates. Whereas cortex viscosity also affects bleb size, with larger blebs observed in a more viscous cortex, the overall change in blebs size and its percentage loss across the range of linker protein stiffness examined appears marginal (Fig. 10B). A similar effect can be observed for cortex permeability in (Fig. 10E).

**Figure 10:**
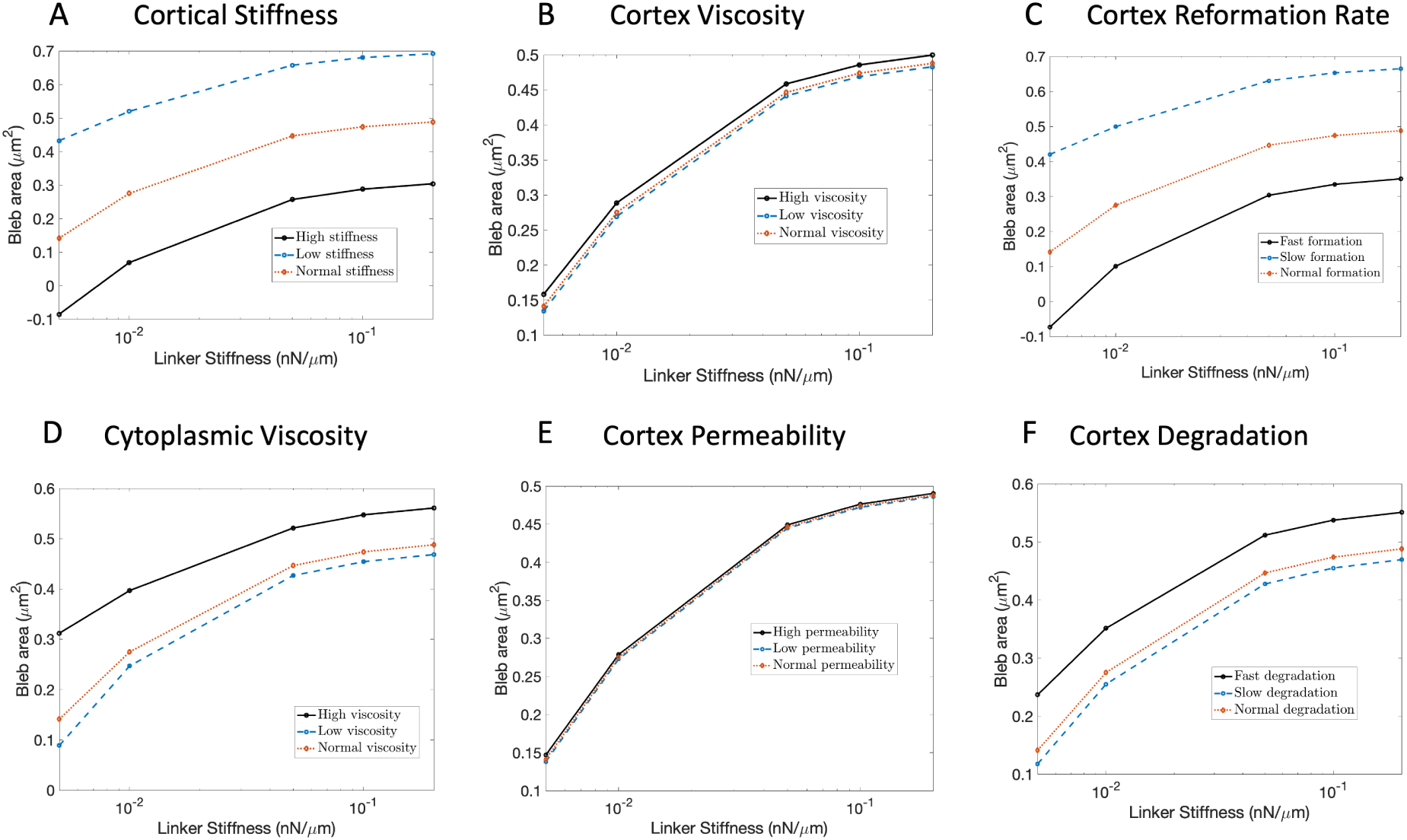
Influence of linker stiffness on bleb size under different viscoelastic conditions. Graphs correspond to bleb size as linker stiffness varies for high (fast), low (slow) and normal elastic or viscous properties of the cell. The normal parameters are those reported in Table 2. High values correspond to twice the normal parameters whereas slow values correspond to half the normal parameters.

## DISCUSSION

Our work investigates the role of TalA and actin dynamics in regulating bleb size and frequency. TalA is known to localize to the posterior of chemotaxing *Dictyostelium* cells and helps to direct blebs to the leading edge of the cell (8, 20). A recent study suggested that TalA enrichment is inversely proportional to blebbing after observing reduced blebbing in regions with a low amount of linker proteins and profusive blebbing in a double mutant (*talA*/*talB* null)(8). We wanted to test whether this phenotype would be observed in a single deletion of *talA*. We find that *Dictyostelium* cells without TalA rarely bleb. These mutant cells also have a larger surface area and produce significantly smaller blebs compared to wild type cells (Fig. 3). This result is startling and in direct contrast to the existing school of thought that cells would increase blebbing in response to an absence of TalA (8). Indeed, one would expect that by removing one of the two known linker proteins in *Dictyostelium*, the attachment of the membrane to the cortex would be significantly reduced, making it easier for blebs to form. Since bleb size correlates with the amoung of intracellular pressure in cells, our observations suggest that *talA* null cells have reduced intracellular pressure, which makes it less likely for blebs to form. Thus, in addition to directing blebs to the front of the cell, this study suggests that TalA helps to maintain the intracellular pressure required for bleb formation.

We hypothesize that the pressure reduction in *talA* null cells comes from a uniform extension of the cell membrane away from the cortex, due to the presence of a much weaker attachment of the membrane to the cortex via say *talB*. To verify the mechanical viability of this hypothesis, we developed a mathematical model of bleb expansion and examined the effect of weakening the strength of linker proteins on bleb size and cell area. Our model improves upon existing models of bleb expansion by allowing for a dynamic change in the density of linker proteins in response to the prevailing hydrostatic pressure, prior to bleb formation. This framework is needed to capture the cells initial effort to balance any excess pressure by deforming the membrane before linker proteins are detached to form blebs. The model also accounts for the dynamic formation of a cortex beneath the protruded membrane as well as its degradation at the old location of the cortex, with parameters determined from a prior study analyzing blebbing data (7). Our model also allows for an investigation of the effect of actin dynamics on bleb formation, something that has not been studied mechanistically hitherto.

Simulations of our model show that decreasing the linker protein strength indeed decreases bleb size (Fig. 9) and causes a greater uniform outward extension of the cell boundary in non-blebbing regions (Fig. 8B). We reason that the extension of the cell boundary increases the surface area to volume ratio of the cell and consequently decreases the prevailing intracellular pressure. This effect is captured by the volume preserving term in our model for pressure (Eq. 16). The key outcome of our modeling effort is in the conclusion that the extension of the cell boundary due to weaker linker proteins, though small (Fig. 8B,C), is sufficient to reduce the intracellular pressure and limit bleb size. This conclusion is not obvious from our experimental results and necessitated mathematical modeling. Additionally, our model simulations and equilibrium analysis support the existing view point that bleb size is regulated by membrane and cortical stiffness as well as intracellular pressure (18, 28), with cortex reformation playing a critical role in determining the steady-state bleb size. We observed that blebs expand to a larger size as the ratio of actin polymerization rate to depolymerization increases in the reforming cortex (Fig. 5D, Fig. 7D). Interestingly, we observed that blebs undergo a partial retraction when the polymerization rate is less than or equal to the depolymerization rate and expand monotonically, otherwise. It is important to note that this partial retraction moves the bleb towards its steady-state size and is not a full withdrawal of the bleb into the cell body as discussed in other studies. Such partial retraction is typical of blebs used for movement, unlike blebs that may be generated by processes such as apoptosis. In particular, Tinevez observed bleb retraction several minutes after bleb stabilization (18), in contrast to our retraction that occurs within the first 20 seconds. In Tinevez’s work the blebs are initiated by laser ablation and hence not motility driven. These blebs also retract fully after the bleb reaches its steady-state size.Hitherto, bleb retraction has been soley associated with the contractile activity of myosin (4, 16, 43). This study suggests that retraction of blebs to their steady-state size can be driven by actin dynamics without the need to invoke myosin II.

In Fig. 10 we examined the effect of various mechanical properties of the cell on the percentage change in blebs due to a reduction in the strength of linker proteins. This analysis is meant to understand how different cell types may respond to weakening the strength of linker proteins. We observed that the stiffness of the cortex as well as its reformation rate beneath the newly formed bleb influence both the size of blebs as well as the percentage reduction in bleb size when the strength of linker proteins is reduced. This result is in agreement with the effect of cortical stiffness and cortex reformation rate on bleb size seen from our 1D and 2D validation results (Fig. 7F,G, Fig. 4C and Fig. 5D), and suggests that the effect of reduced membrane-cortex linker strength on bleb size is exacerbated in cells with stiffer cortex as well as those that reform the cortex rapidly. This conclusion makes sense since cells with higher cortical stiffness will tend to limit the expansion of the membrane once cortex reformation begins. This is in spite of the fact that a stiffer cortex will limit the initial uniform expansion of the membrane in response to the application of excess pressure, making more intracellular pressure available for bleb formation (Fig. S2A), whereas the cortex reformation rate will have no effect on the intracellular pressure available for bleb formation (Fig. S2C).

Surprisingly, our results Fig. 10 revealed an effect of cytoplasmic viscosity and cortex viscosity on bleb size as well as the percentage reduction in blebs due to weakening linker proteins. In particular, we found that reducing the strength of linker proteins in cells with lower cytoplasmic viscosity or cortex viscosity results in a greater reduction in bleb size, with the effect of cortex viscosity being marginal. These results appear to be in stark contrast to the validation study in our 1D model which predicted that the viscous properties of the cell only increase the speed of bleb expansion but have no affect the steady-state bleb size (Fig. 4D, Fig. 5A). This claim was also supported by the absence of these viscous parameters in the calculation of 1) the critical density of linker proteins and displacement of the cell boundary prior to bleb initiation (see Eq. 11) and 2) the final displacement of the cell boundary after bleb initiation (see Eqn. 18).

Recall that the primary objective for developing the 2D model was to create a more accurate representation of the pressure and boundary dynamics of blebbing cells, which could not be accounted for within a 1D framework. One of the primary limitations of our 1D model is the use of a phenomenological pressure term that computes the driving force for blebbing based solely on the local membrane displacement from the cortex. Additionally, in computing the initial driving pressure for bleb expansion, we only took into consideration the critical displacement of the cell boundary at the point where linker proteins are detached, completely ignoring the boundary dynamics in other locations. These simplifying assumptions appear to have made a significant difference in the role of the viscous components of cell on steady-state bleb size. Our results in Fig. 8 demonstrate that, in locations where blebs do not form, the cell boundary experiences a small uniform expansion in response to the excess pressure driving bleb expansion. This boundary expansion in non-blebbing regions is reduced by increasing the viscosity of the cytoplasm (see Fig. S2D), thereby making more pressure available to form larger blebs as seen in Fig. 10D . The effect of cortex viscosity on the expansion of the boundary in non-blebbing regions is limited (see Fig. S2B), thus explaining its marginal effect on bleb size observed in Fig. 10B. Other factors such as cortex permeability, cortex degradation rate and cortex reformation rate were observed to have no effect on boundary expansion in non-blebbing regions (see Fig. S2 C,E,F).

For simplicity, our model for bleb expansion assumes a viscous cytoplasm with a uniform distribution of intracellular pressure. This is limiting, since other studies have pointed to the cytoplasm being poroelastic and permitting short-lived pressure gradients within the cell.Our modeling and experiments suggest that weakening the strength of linker proteins results in a uniform extension of the membrane from the cortex, which limits the driving pressure for bleb formation. Consequently, a double mutant should have a more significant reduction in bleb size (since there will be more pressure relief) and be less likely to make blebs. It is unclear to us why a double deletion of *talA* and *talB* would result in profusive blebbing, as suggested in (8). One possibility is that these proteins have very different roles in blebbing and as such more experiments need to be done to quantify blebbing in a single *talB* deletion. The experimental data presented in (8) to support profusive blebbing of the double *talA,talB* mutant shows membrane protrusions without visible cytoplasm. It is unclear if these go through the bleb cycle described in Fig. 2. There is also no quantification of the frequency or size of blebs in comparison to wild type. Thus, it will be informative to conduct new experiments for the double mutant to clarify the role of *talB* in blebbing.

## AUTHOR CONTRIBUTIONS

S.H.D.S. and E.A.A developed, analyzed, simulated mathematical model and wrote manuscript. E.M and J.R analyzed experimental data. Z.S performed under-agarose experiments. D.B supervised experiments and edited article.

## ACKNOWLEDGEMENTS

This work was supported by grants to DB from the National Science Foundation (MCB-1244162), a PSC-CUNY grant (692710047), as well as Research Centers in Minority Institutions Program grants from the National Institute on Minority Health and Health Disparities (8 G12 MD007599) from the National Institutes of Health. ZS was supported by the Research Initiative for Scientific Enhancement (RISE) program at Hunter College which is funded by the NIH (GM060665, https://www.nih.gov).

