## Supplementary Information for "A mathematical model for bleb expansion clarifies a role for TalA in regulating blebbing"

### S1 Supplemental figures

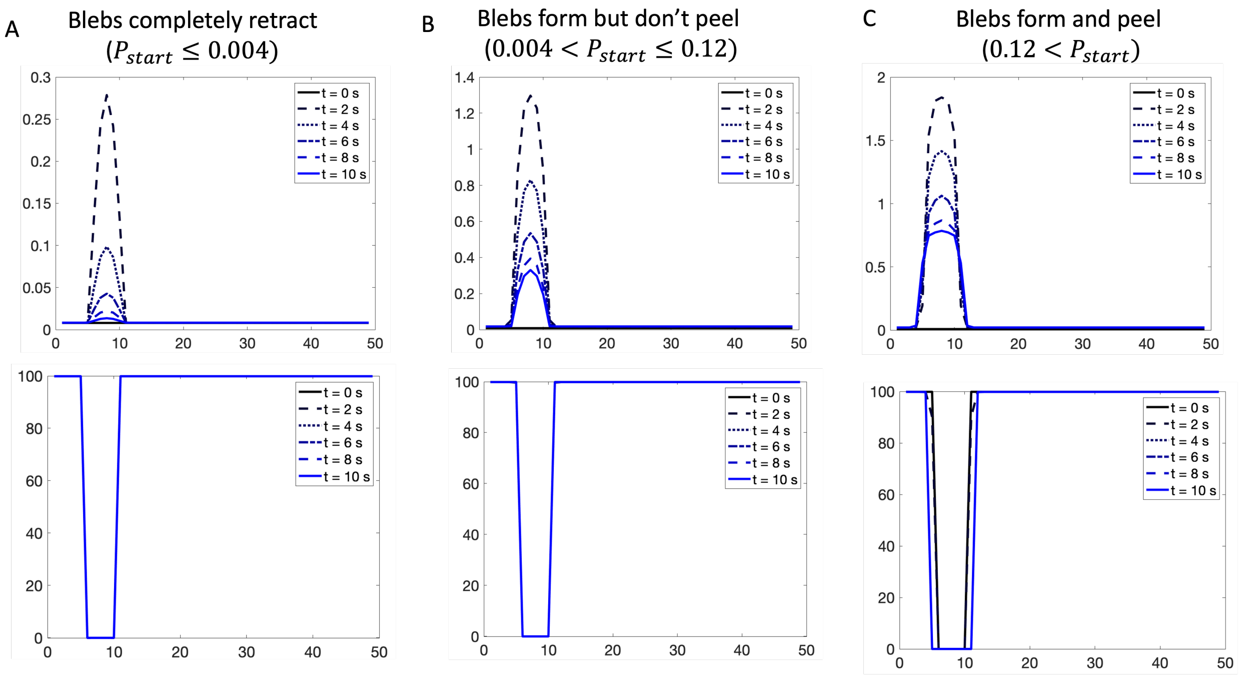


Figure S1: **Conditions for bleb formation and peeling.** Range of excess pressure for which A) blebs fail to form, B) blebs form without peeling, C) blebs form and peel. Peeling factor is set to 6 for all cases.

### S2 Supplemental methods

##### Derivation of time-dependent mechanical parameters

Our primary concern is to obtain an expression for $r_1(t)$ and $r_2(t)$ used to formulate the time dependent mechanical parameter $\tau_y(t), \tau_c(t)$ and $k_b(t)$ of the 1D model. Let $a_{bleb}(t)$ and $a_{scar}(t)$ denote the actin concentration in the developing and old cortex at any time and $a_0$ the concentration of actin the mature cortex. Then we set $r_1(t) = \frac{a_{bleb}(t)}{a_0}$ and $r_2(t) = \left(1 - \frac{a_{scar}(t)}{a_0}\right)$. The expression for $r_1(t)$ is precisely the fraction of actin in the developing cortex at any time. The expression for $r_2(t)$ represents the fraction of actin lost in the degrading cortex, which correlates positively with the viscosity of the fluid within the bleb. It remains to obtain expressions for the actin concentration in the bleb cortex $a_{bleb}(t)$ and actin scar $a_{scar}(t)$.

Recently, we introduced the following linear model for actin a(t) and myosin m(t) dynamics during reformation of the bleb cortex and degradation of the actin scar [1],

$$\begin{aligned} \frac{da}{dt} &= k_a^{on} - k_a^{off} a(t), \\ \frac{dm}{dt} &= k_m^{on} a(t) - k_m^{off} m(t). \end{aligned} \tag{S1}$$

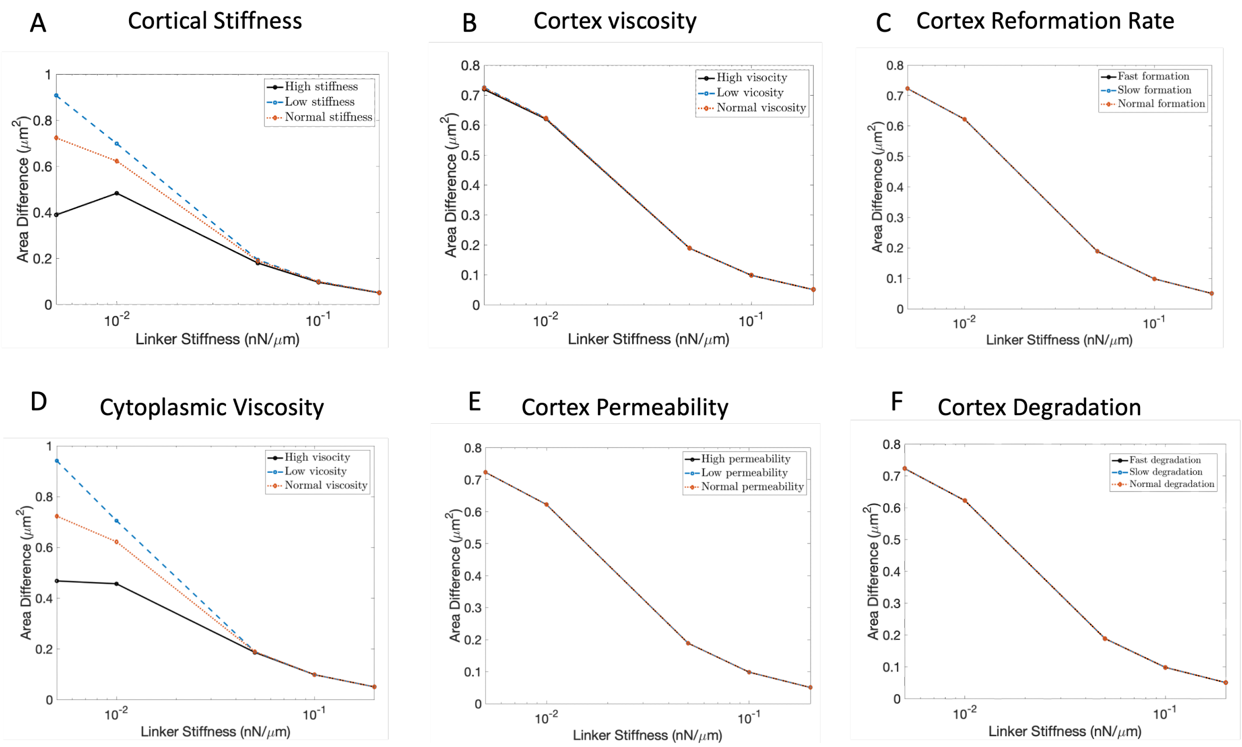


Figure S2: **Effect of cell mechanical properties on uniform expansion of cell without bleb initiation.** Graphs correspond to cell area difference in response to excess pressure as linker stiffness varies for high (fast), low (slow) and normal elastic or viscous properties of the cell. The normal parameters are those reported in the main manuscript. High values correspond to twice the normal parameters whereas slow values correspond to half the normal parameters

The model parameters were estimated using blebbing data from *Dictyostelium discoideum* cells migrating under the same experimental conditions used to obtain the data presented in this work. Here, $k_a^{on}, k_a^{off}$ describe the polymerization and depolymerization rates of actin in the cortex. The estimated model parameters differ between reformation of the bleb cortex and degradation of the actin scar. Hence, we will denote the respective rates at the bleb cortex by $k_{ab}^{on}, k_{ab}^{off}$ and those at the actin scar by $k_{as}^{on}, k_{as}^{off}$. Whereas this model fits the experimental data on actin and myosin concentration in the reforming bleb cortex well, it was only able to capture the major trends of actin and myosin concentration in the degrading actin scar [1]. Nevertheless, the simplicity in the decoupling of actin dynamics from myosin make this an attractive model for estimating the relative concentration of actin in the bleb cortex and actin scar.

Solving for the actin concentration $a(t)$ from Eq. S1 we obtain

$$a(t) = \frac{k_{ab}^{on}}{k_{ab}^{off}} + \left( a(0) - \frac{k_{ab}^{on}}{k_{ab}^{off}} \right) e^{-k_{ab}^{off} t}$$

where $a(0)$ is the initial density of actin in the reforming cortex and $\frac{k_{ab}^{on}}{k_{ab}^{off}}$ is the equilibrium density of actin (referred to elsewhere as $a_{rest}$). At the reforming bleb cortex $a(0) = 0$, thus the relative actin density there is given by

$$a_{bleb}(t) = \frac{k_{ab}^{on}}{k_{ab}^{off}}(1 - e^{-k_{ab}^{off} t}). \tag{S2}$$

At the degrading actin scar $a(0) = a_0 \neq 0$. Our estimated value for $k_{as}^{on}$ was near zero in [1]. Setting $k_{as}^{on} = 0$, we obtain the actin density

$$a_{scar}(t) = a_0 e^{-k_{as}^{off} t}$$

in the degrading actin scar.

#### S2.1 Steady-state analysis

The motivation for this section is two fold. First, we will determine the critical displacement $u^0$ and linker density $\rho_a^0$ necessary for initializing bleb expansion in the 1D model. Since bleb expansion is expected to stall over time, we will follow this analysis with an investigation of the parameters that control the steady-state bleb size.

##### Critical displacement and linker density

The initial conditions for our 1D bleb expansion model are the critical displacement $u^0$ and critical linker density $\rho_a^0$.To calculate these values, we set $\dot{u}(t) = 0$ and $\dot{\rho}_a(t) = 0$ in the 1D bleb expansion model

$$\frac{du}{dt} = \frac{F_{1D}}{(\tau_c + \tau_y)} - \frac{(k_b + k_a \rho_a)}{(\tau_c + \tau_y)} u \tag{S3}$$

$$\frac{d\rho_a}{dt} = k_{on}(\rho_0 - \rho_a) - k_{off}(u(t))\rho_a \tag{S4}$$

with all mechanical parameters fixed. This yields

$$\frac{F_{1D}}{\tau_c + \tau_y} - \frac{\left(k_b + k_a \rho_a^0\right)}{\tau_c + \tau_y} u^0 = 0 \tag{S5}$$

$$k_{on}\left[\rho_0 - \rho_a^0\right] - k_{off}(u^0)\rho_a^0 = 0. \tag{S6}$$

After substituting our driving force $F_{1D} = F_0 e^{-mu}$, our equilibrium solutions thus satisfy the following nonlinear system of equations

$$u^0 = \frac{e^{-mu^0}}{\left(k_b + k_a \rho_a^0\right)} \tag{S7}$$

$$\rho_a^0 = \frac{k_{on}\rho_0}{k_{on} + k_{off}^0 e^{\delta \beta k_a u^0}}. \tag{S8}$$

##### Steady-state bleb size

Recall that once the membrane detaches from the cortex, linker proteins are lost and no longer contribute to bleb dynamics. Hence, we set $\rho_a = 0$ and ignore its dynamics in the 1D model (Eq. S3). Our main equation for studying equilibrium bleb size is thus,

$$\dot{u} = \frac{F_{1D}(u)}{\tau_c(t) + \tau_y(t)} - \frac{k_b(t)}{\tau_c(t) + \tau_y(t)} u(t) \tag{S9}$$

which is non-autonomous. First we convert it to an autonomous system by using the exact forms of $k_b(t), \tau_c(t)$ and $\tau_y(t)$ given in the main paper and their rates of change

$$\begin{aligned}\frac{dk_b}{dt} &= k_c \frac{\dot{a}_{bleb}(t)}{a_0}\\ \frac{d\tau_c}{dt} &= \tau_c^0 \frac{\dot{a}_{bleb}(t)}{a_0}\\ \frac{d\tau_y}{dt} &= -(1-\theta)\tau_y^0 \dot{a}_{scar}.\end{aligned} \tag{S10}$$

Recall from Eq. S1 that

$$\begin{aligned}\dot{a}_{bleb} &= k_{ab}^{on} - k_{ab}^{off} a_{bleb}\\ \dot{a}_{scar} &= -k_{as}^{off} a_{scar}.\end{aligned}$$

Therefore, the complete autonomous system becomes,

$$\begin{aligned}\frac{du}{dt} &= \frac{F_{1D}(u)}{\tau_c+\tau_y} - \frac{k_b}{\tau_c+\tau_y} u\\ \frac{da_{bleb}}{dt} &= k_{ab}^{on} - k_{ab}^{off} a_{bleb}\\ \frac{da_{scar}}{dt} &= -k_{as}^{off} a_{scar}\\ \frac{dk_b}{dt} &= k_c \frac{(k_{ab}^{on} - k_{ab}^{off} a_{bleb})}{a_0}\\ \frac{d\tau_c}{dt} &= \tau_c^0 \frac{(k_{ab}^{on} - k_{ab}^{off} a_{bleb})}{a_0}\\ \frac{d\tau_y}{dt} &= (1-\theta)\tau_y^0 k_{as}^{off} a_{scar}.\end{aligned} \tag{S11}$$

#### Steady state/equilibrium solution

At equilibrium, we have

$$\frac{du}{dt} = \frac{dk_b}{dt} = \frac{d\tau_c}{dt} = \frac{d\tau_y}{dt} = \frac{da_{bleb}}{dt} = \frac{da_{scar}}{dt} = 0,$$

resulting in the system of equations

$$\frac{F_{1D}(u)}{\tau_c+\tau_y} - \frac{k_b}{\tau_c+\tau_y} u = 0 \tag{S12}$$

$$k_{ab}^{on} - k_{ab}^{off} a_{bleb} = 0 \tag{S13}$$

$$-k_{as}^{off} a_{scar} = 0 \tag{S14}$$

$$k_c \frac{(k_{ab}^{on} - k_{ab}^{off} a_{bleb})}{a_0} = 0 \tag{S15}$$

$$\tau_c^0 \frac{(k_{ab}^{on} - k_{ab}^{off} a_{bleb})}{a_0} = 0 \tag{S16}$$

$$(1-\theta)\tau_y^0 k_{as}^{off} a_{scar} = 0.. \tag{S17}$$

We denote the equilibrium value of an arbitrary variable $w$ as $w^*$. Note from Eq. S14 that $a^*_{scar} = 0$, which implies that Eq. S17 is trivially satisfied. From Eq. S13, $a^*_{bleb} = \frac{k^{on}_{ab}}{k^{off}_{ab}}$, which also implies that Eq. S16 and Eq. S15 are trivially satisfied. By allowing $t \to \infty$, we find that $\tau^*_y = \tau^0_y, \tau^*_c = \frac{\tau^0_c k^{on}_{ab}}{a_0 k^{off}_{ab}}$ and $k^*_b = k_m + \frac{k_c k^{on}_{ab}}{a_0 k^{off}_{ab}}$. Therefore, from Eqn. S12,

$$F_{1D}(u^*) = k^*_b u^*.$$

Using the model for $F(u) = F_0 e^{-mu}$, the equilibrium bleb size, $u^*$ can be obtained by solving the nonlinear equation

$$F_0 e^{-mu^*} - \frac{k_c k^{on}_{ab}}{a_0 k^{off}_{ab}} u^* = k_m. \tag{S18}$$

#### S2.2 2D model of bleb expansion

##### Submodel for displacement and linker density

At a fixed point $(x(s,t), y(s,t))$ on the cell boundary, the 1D velocity is,

$$\dot{u}(t) = \frac{F_{1D}}{\tau_c(t) + \tau_y(t)} - \frac{\big(k_b(t) + k_a \rho_a(t)\big)}{\tau_c(t) + \tau_y(t)} u(t).$$

Setting $\rho_a(t) = \Phi(x, y, t)$, we define the 2D velocity as

$$\frac{dI}{dt} = \frac{F_{2D}\hat{n}}{\tau_c(s,t) + \tau_y(s,t)} - \frac{\big(k_b(s,t) + k_a \Phi(x,y,t)\big)}{\tau_c(s,t) + \tau_y(s,t)} I. \tag{S19}$$

where

$$\tau_y(s,t) = \tau^0_y(1 - X_{bleb}(s)) + [\theta\tau^0_y + (1-\theta)\tau^0_y(1 - e^{-k^{off}_{as} t})]X_{bleb}(s) \tag{S20}$$

$$\tau_c(s,t) = \tau^0_c(1 - X_{bleb}(s)) + [\frac{\tau^0_c k^{on}_{ab}(1 - e^{-k^{off}_{ab} t})}{a_0 k^{off}_{ab}}]X_{bleb}(s) \tag{S21}$$

$$k_b(s,t) = (k_m + k_c)(1 - X_{bleb}(s)) + \left[k_m + \frac{k_c k^{on}_{ab}(1 - e^{-k^{off}_{ab} t})}{a_0 k^{off}_{ab}}\right] X_{bleb}(s). \tag{S22}$$

Suppose the displacement vector $I$ and linker density $\Phi$ are defined off the cell boundary, we observe that

$$\begin{aligned}\frac{dI(x,y,t)}{dt} &= \frac{\partial I}{\partial x}\frac{dx}{dt} + \frac{\partial I}{\partial y}\frac{dy}{dt} + \frac{\partial I}{\partial t} \\ &= DI\vec{v} + \frac{\partial I}{\partial t}\end{aligned} \tag{S23}$$

where $\vec{v} = \left(\frac{dx}{dt}, \frac{dy}{dt}\right)$ and $DI$ is the Jacobian matrix of the displacement vector $I = (I_1, I_2)$. Substituting Eq. S23 into Eq. S19 and expanding out the components of the displacement vector, we obtain the following model for the 2D velocity

$$\frac{\partial I_1}{\partial t} + \nabla I_1.\vec{v} = \frac{F_{2D}\hat{n}_x}{\tau_c(s,t)+\tau_y(s,t)} - \frac{\big(k_b(s,t)+k_a\Phi(x,y,t)\big)}{\tau_c(s,t)+\tau_y(s,t)} I_1 \tag{S24}$$

$$\frac{\partial I_2}{\partial t} + \nabla I_2.\vec{v} = \frac{F_{2D}\hat{n}_y}{\tau_c(s,t)+\tau_y(s,t)} - \frac{\big(k_b(s,t)+k_a\Phi(x,y,t)\big)}{\tau_c(s,t)+\tau_y(s,t)} I_2 \tag{S25}$$

Similarly, using the 1D model for linker protein density in the main text with $\rho_a = \Phi$ and $|I| = u$, we can write the linker protein density dynamics in 2D as

$$\frac{\partial \Phi}{\partial t} + \nabla \Phi.\vec{v} = k_{on}[\rho_0 - \Phi] - k_{off}(|I|)\Phi. \tag{S26}$$

The gradient terms in Eq. S24, Eq. S25 and Eq. S26 assume that the displacement vector and linker density are defined off the cell boundary. However, these quantities only have physical significance on the boundary of the cell. To resolve this, we extend them off the boundary by assuming that $I, \Phi$ are constant in the normal direction and consequently impose the constraints

$$\begin{aligned} \nabla I_i \cdot \hat{n} &= 0, \qquad i = 1,2 \\ \nabla \Phi \cdot \hat{n} &= 0. \end{aligned} \tag{S27}$$

We now proceed to derive expressions for $\nabla I_i \cdot \vec{v}$ and $\nabla \Phi \cdot \vec{v}$ that only depend on derivatives taken along the cell boundary. This framework where derivatives are expressed in terms of the spatial parameter around the cell boundary is new to this work.

Let $f(x(s), y(s))$ be some scalar quantity defined along the cell boundary with unit tangent vector $\hat{\tau}$ and unit outward normal vector $\hat{n}$ at location $(x(s), y(s))$. Then,

$$\nabla f = (\nabla f \cdot \hat{\tau})\hat{\tau} + (\nabla f \cdot \hat{n})\hat{n}.$$

Under the assumption that $\nabla f \cdot \hat{n} = 0$ the gradient is reduced to its tangential component. Recall that,

$$\begin{aligned} \frac{d}{ds} f(x(s), y(s)) &= \frac{\partial f}{\partial x}\frac{dx}{ds} + \frac{\partial f}{\partial y}\frac{dy}{ds} \\ &= \nabla f . \frac{d\vec{x}}{ds}, \end{aligned}$$

where $\vec{x} = (x(s), y(s))$ and $\vec{\tau} = \frac{d\vec{x}}{ds}$ is the tangent vector at the point $(x(s), y(s))$. Dividing both sides by $|\vec{\tau}|$ we obtain

$$\frac{1}{|\vec{\tau}|}\frac{df}{ds} = \nabla f \cdot \hat{\tau},$$

which after multiplying with $\hat{\tau}$ results in

$$\frac{\vec{\tau}}{|\vec{\tau}|^2}\frac{df}{ds} = (\nabla f \cdot \hat{\tau})\hat{\tau} = \nabla f. \tag{S28}$$

With the expression for the gradient above we can write

$$\nabla I_1 \cdot \vec{v} = \frac{\vec{\tau} \cdot \vec{v}}{|\vec{\tau}|^2} \frac{dI_1}{ds} \tag{S29}$$

$$\nabla I_2 \cdot \vec{v} = \frac{\vec{\tau} \cdot \vec{v}}{|\vec{\tau}|^2} \frac{dI_2}{ds} \tag{S30}$$

$$\nabla \Phi \cdot \vec{v} = \frac{\vec{\tau} \cdot \vec{v}}{|\vec{\tau}|^2} \frac{d\Phi}{ds}. \tag{S31}$$

Now we set, $\vec{v} = \alpha \frac{dI}{dt}$ for $\alpha \geq 1$ by assuming the velocity to be proportional to $\frac{dI}{dt}$. Substituting the expression for $\frac{dI}{dt}$ given in Eq. S19 into Eq. S29- Eq. S31, we obtain

$$\nabla I_1 \cdot \vec{v} = -\alpha \frac{(\vec{\tau} \cdot I)(k_b + k_a P)}{|\vec{\tau}|^2(\tau_c + \tau_y)} \frac{dI_1}{ds} \tag{S32}$$

$$\nabla I_2 \cdot \vec{v} = -\alpha \frac{(\vec{\tau} \cdot I)(k_b + k_a P)}{|\vec{\tau}|^2(\tau_c + \tau_y)} \frac{dI_2}{ds} \tag{S33}$$

$$\nabla \Phi \cdot \vec{v} = -\alpha \frac{(\vec{\tau} \cdot I)(k_b + k_a \Phi)}{|\vec{\tau}|^2(\tau_c + \tau_y)} \frac{d\Phi}{ds}. \tag{S34}$$

Now, substituting (Eq. S32-Eq. S34) into the model equations in Eq. S24- Eq. S26 and using the fact that $I = (I_1, I_2)$ we obtain the following model for bleb expansion in 2D

$$\frac{\partial I}{\partial t} - \alpha \frac{\vec{\tau}.I}{|\vec{\tau}|^2} \frac{(k_b + k_a \Phi)}{\tau_c + \tau_y} \frac{\partial I}{\partial s} = \frac{F_{2D}\hat{n}}{\tau_c + \tau_y} - \frac{\left(k_b + k_a \Phi\right)}{\tau_c + \tau_y} I \tag{S35}$$

$$\frac{\partial \Phi}{\partial t} - \alpha \frac{\vec{\tau}.I}{|\vec{\tau}|^2} \frac{(k_b + k_a \Phi)}{\tau_c + \tau_y} \frac{\partial \Phi}{\partial s} = k_{on}[\rho_0 - \Phi] - k_{off}(|I|)\Phi \tag{S36}$$

where

$$\begin{aligned}
\tau_y(s,t) &= \tau_y^0(1 - X_{bleb}(s)) + [\theta \tau_y^0 + (1-\theta)\tau_y^0(1 - e^{-k_{as}^{off} t})]X_{bleb}(s) \\
\tau_c(s,t) &= \tau_c^0(1 - X_{bleb}(s)) + [\frac{\tau_c^0 k_{ab}^{on}(1 - e^{-k_{ab}^{off} t})}{a_0 k_{ab}^{off}}]X_{bleb}(s) \\
k_b(s,t) &= (k_m + k_c)(1 - X_{bleb}(s)) + \left[k_m + k_c \frac{k_{ab}^{on}(1 - e^{-k_{ab}^{off} t})}{a_0 k_{ab}^{off}}\right] X_{bleb}(s).
\end{aligned} \tag{S37}$$

#### S2.3 Numerical solution of 2D blebbing model

##### Reformulation of PDE as ODE system

We simulate the model Eq. S35- Eq. S37, using the particle tracking approach. Here, discrete points along the cell boundary are evolved in time using the proposed mathematical model. Given the cell boundary $\Gamma = \{(x(s), y(s)) : s \in [a,b]\}$, define $s_i = a + ih, \quad i = 0, 1, 2, \ldots, m$ and $h = \frac{b-a}{m}$ and set $(x(s_i), y(s_i)) = (x_i, y_i)$. Then the model will be discretized on the partition $\Lambda_h := \{(x_i, y_i)\}_{i=0}^m$.

To simplify the notation, set

$$a(I,\Phi,s,t) = \alpha\frac{\vec{\tau}.I}{|\vec{\tau}|^2}\frac{(k_b(s,t)+k_a\Phi(s,t))}{\tau_c(s,t)+\tau_y(s,t)}$$
$$b(I,\Phi,s,t) = \frac{F_{2D}(s,t)\hat{n}}{\tau_c(s,t)+\tau_y(s,t)} - \frac{\left(k_b(s,t)+k_a\Phi(s,t)\right)}{\tau_c(s,t)+\tau_y(s,t)}I$$
$$c(I,\Phi) = k_{on}[\rho_0-\Phi] - k_{off}(|I|)\Phi.$$

Then the model reduces to the form

$$\begin{aligned}\frac{\partial I}{\partial t} &= a(I,\Phi,s,t)\frac{\partial I}{\partial s} + b(I,\Phi,s,t)\\ \frac{\partial \Phi}{\partial t} &= a(I,\Phi,s,t)\frac{\partial \Phi}{\partial s} + c(I,\Phi).\end{aligned} \tag{S38}$$

Simplifying further, by writing as a single equation, set $W = (I,\Phi)^T$ and $f(W,s,t) = (b(I,\Phi,s,t), c(I,\Phi))^T$. Then, our model becomes,

$$\frac{\partial W}{\partial t} = a(W,s,t)\frac{\partial W}{\partial s} + f(W,s,t). \tag{S39}$$

Next, at the location $(x_i, y_i)$, the spatial derivatives of $W$ are discritized with the second order central difference formula,

$$\frac{\partial W}{\partial s}|_{s_i} \approx \frac{W_{i+1}-W_{i-1}}{2h}$$

yielding the spatially discrete equations

$$\frac{dW_i}{dt} = a(W_i,s_i,t)\left(\frac{W_{i+1}-W_{i-1}}{2h}\right) + f(W_i,s_i,t), \quad i = 0,1,2,\ldots,m. \tag{S40}$$

Now, since the boundary of the cell is closed, we can set

$$W_{m+1} = W_0; W_{-1} = W_m$$

and rewrite the model equations as the ode system

$$\frac{dW_h}{dt} = G(W_h, s_h, t) \tag{S41}$$

where, $W_h = (W_0, W_1, W_2, \cdots, W_m)^T$, $s_h = (s_0, s_1, s_2, \cdots, s_m)^T$ and

$$G(W_h,s_h,t) = \begin{pmatrix} a(W_0,s_0,t)\left(\frac{W_1-W_m}{2h}\right) + f(W_0,s_0,t)\\ a(W_1,s_1,t)\left(\frac{W_2-W_0}{2h}\right) + f(W_1,s_1,t)\\ a(W_2,s_2,t)\left(\frac{W_3-W_1}{2h}\right) + f(W_2,s_2,t)\\ \vdots\\ a(W_{m-1},s_{m-1},t)\left(\frac{W_m-W_{m-2}}{2h}\right) + f(W_{m-1},s_{m-1},t)\\ a(W_m,s_m,t)\left(\frac{W_0-W_{m-1}}{2h}\right) + f(W_m,s_m,t). \end{pmatrix} \tag{S42}$$

The ode system in Eq. S41 is solved in MATLAB using ode113 with initial conditions

$$(W_h(0))_i := (I_i(0), \Phi_i(0)) = (u^0\hat{n}_i, \rho^0),$$

where $u^0$ and $\rho^0$ are the critical strain and linker density described in Eq. S7 and Eq. S8 and $\hat{n}_i$ is the unit normal vector to the cell boundary at the point $(x_i, y_i)$. In the discussion that follows, we describe the estimation of normal and tangent vectors.

#### Computing curvature, normal and tangent vectors

The accuracy of our model relies heavily on the correct calculation of the intracellular pressure and the direction of bleb expansion, all of which depend on the boundary curvature, normal and tangent vectors. Consider a closed curve parameterized as $(x(s), y(s))$, then the curvature is defined as

$$\frac{|x^{'}y^{''} - x^{''}y^{'}|}{((x^{'})^2 + (y^{'})^2)^{3/2}}$$

with unit tangent and normal vectors given by

$$\hat{\tau} = (x', y')^T \frac{1}{\sqrt{(x^{'})^2 + (y^{'})^2}}, \quad \hat{n} = (y^{'}, -x^{'})^T \frac{1}{\sqrt{(x^{'})^2 + (y^{'})^2}}.$$

To ensure that curvature, normal and tangent vectors are defined continuously, we approximated the cell boundary with a parametric cubic spline, at each time step. The cubic spline was constructed using the 'cscvn' function in MATLAB and evaluated, along with its derivatives, on a given partition using the 'fnder' and 'fnval' functions in MATLAB. At the shoulder points of the bleb, where the bleb boundary meets the cell boundary, we defined the normal and tangent vectors by averaging the corresponding vectors computed from the two boundary profiles.

#### Calculating cell area

Recall from Green's Theorem, that the area of the region $D$ bounded by a positively oriented closed curve $C$ with parameterization $(x(s), y(s))$, is given by

$$\int_C x \; dy = \iint_D dA$$

For our purposes, $C$ is the cell boundary $\Gamma$ and $D$ its area. Consider the following parametric cubic spline interpolation of the boundary partition $\Lambda_h$, with equations for the the $i^{th}$ segment given as

$$x_i(s) = a_{ix} + b_{ix}s + c_{ix}s^2 + d_{ix}s^3$$
$$y_i(s) = a_{iy} + b_{iy}s + c_{iy}s^2 + d_{iy}s^3$$

$i = 1, 2, \ldots, m$. Then,

$$
\begin{aligned}
\int_{\Gamma} x\ dy &= \int_{\Gamma} xy^{'}(s)\ ds \\
&= \sum_{i=1}^{m} \int_{s_{i-1}}^{s_i} (a_{ix} + b_{ix}s + c_{ix}s^2 + d_{ix}s^3).(b_{iy} + 2c_{iy}s + 3d_{iy}s^2)ds \\
&= \sum_{i=1}^{m} \int_{s_{i-1}}^{s_i} [(a_{ix}b_{iy}) + (2a_{ix}c_{iy} + b_{ix}b_{iy})s + (3a_{ix}d_{iy} + 2b_{ix}c_{iy} + c_{ix}b_{iy})s^2 + \\
&(3b_{ix}d_{iy} + 2c_{ix}c_{iy} + d_{ix}b_{iy})s^3 + (3c_{ix}d_{iy} + 2d_{ix}c_{iy})s^4 + 3d_{ix}d_{iy}s^5]ds \\
&= \sum_{i=1}^{m} \Bigg[\Bigg((a_{ix}b_{iy})[s_i - s_{i-1}] + (2a_{ix}c_{iy} + b_{ix}b_{iy})\frac{[s_i - s_{i-1}]^2}{2} + \\
&(3a_{ix}d_{iy} + 2b_{ix}c_{iy} + c_{ix}b_{iy})\frac{[s_i - s_{i-1}]^3}{3} + (3b_{ix}d_{iy} + 2c_{ix}c_{iy} + d_{ix}b_{iy})\frac{[s_i - s_{i-1}]^4}{4} + \\
&(3c_{ix}d_{iy} + 2d_{ix}c_{iy})\frac{[s_i - s_{i-1}]^5}{5} + 3d_{ix}d_{iy}\frac{[s_i - s_{i-1}]^6}{6}\Bigg)\Bigg] \\
&= \sum_{i=1}^{m} \Bigg[\Bigg((a_{ix}b_{iy})h + (2a_{ix}c_{iy} + b_{ix}b_{iy})\frac{h^2}{2} + \\
&(3a_{ix}d_{iy} + 2b_{ix}c_{iy} + c_{ix}b_{iy})\frac{h^3}{3} + (3b_{ix}d_{iy} + 2c_{ix}c_{iy} + d_{ix}b_{iy})\frac{h^4}{4} + \\
&(3c_{ix}d_{iy} + 2d_{ix}c_{iy})\frac{h^5}{5} + 3d_{ix}d_{iy}\frac{h^6}{6}\Bigg)\Bigg]
\end{aligned}
$$
